# The piRNA protein Asz1 is essential for germ cell and gonad development in zebrafish and exhibits differential necessities in distinct types of RNP granules

**DOI:** 10.1101/2023.07.18.549483

**Authors:** Adam Ahmad, Yoel Bogoch, Gal Shvaizer, Yaniv M. Elkouby

**Affiliations:** Department of Developmental Biology and Cancer Research, The Hebrew University of Jerusalem Faculty of Medicine, Ein- Kerem Campus, Israel 9112102; Institute for Medical Research – Israel-Canada (IMRIC), Ein- Kerem Campus, Israel 9112102

## Abstract

Germ cells are essential for fertility, embryogenesis, and reproduction. Germline development requires distinct types of RNA-protein (RNP) granules, including germ plasm in embryos, piRNA granules in gonadal germ cells, and the Balbiani body (Bb) in oocytes. However, the regulation of RNP assemblies in zebrafish germline development are still poorly understood. Asz1 is a piRNA protein in Drosophila and mice. Zebrafish Asz1 localizes to both piRNA and Bb granules, with yet unknown functions. Here, we hypothesized that Asz1 functions in RNP granule assemblies and germline development in zebrafish. We generated *asz1* mutant fish to determine the roles of Asz1 in germ cell development. We show that Asz1 is dispensable for somatic development, but essential for germ cell and gonad development. *asz1^-/-^* fish developed exclusively as sterile males with severely underdeveloped testes that lacked germ cells, demonstrating that Asz1 is essential for spermatogenesis. Mechanistically, we provide evidence to conclude that zygotic Asz1 is not required for primordial germ cell specification or migration to the gonad, but is essential for germ cell survival during post-embryonic gonad development, likely by suppressing the expression of germline transposons. Increased transposon expression and morphologically mis-organized piRNA granules in *asz1* mutants, argues that zebrafish Asz1 functions in the piRNA pathway. We generated *asz1;tp53* fish to partially rescue ovarian development, revealing underdeveloped mutant ovaries with defective oocytes, and that Asz1 is also essential for oogenesis. We further showed that in contrast with piRNA granules, Asz1 is dispensable for Bb granule formation, as shown by normal Bb localization of Buc and *dazl*. By uncovering Asz1 as an essential regulator of germ cell survival and gonadogenesis in zebrafish, and determining its differential necessity in distinct RNP granule types, our work advances our understanding of the developmental genetics of reproduction and fertility, as well as of RNP granule biology.

**Author Summary:** Germ cells undergo a highly dynamic developmental program that begins in the early embryo and continues through juvenile and adult life. Identifying functional regulators and deciphering the developmental mechanisms of germ cells are critical for advancing our understanding of fertility and reproduction, as well as their associated diseases. Here, we identified Asz1 as an essential regulator of germ cell and gonad development in zebrafish. We demonstrate that zygotic Asz1 is dispensable for the specification and migration of primordial germ cells in the embryo, but is necessary for germ cell survival in the developing gonad, likely by protecting them from transposable elements. Upon loss of asz1, expression of germline transposons was induced, and piRNA granules were mis-organized, suggesting a conserved role for zebrafish Asz1 in the piRNA pathway. We show that Asz1 is required for both spermatogenesis and oogenesis. However, unlike RNA-protein (RNP) granules of the piRNA pathway, Asz1 was not required for RNP granules of the Balbiani body in differentiating oocytes, revealing its differential necessity in distinct types of germline RNP granules. In mice, Asz1 was shown to be essential for spermatogenesis but not oogenesis, but its functions in human gonads are unclear. Our work reports the functional requirements of Asz1 in both sexes in zebrafish, contributes to our knowledge of developmental reproduction biology, and sheds new light on the complexity of RNP granule assemblies.

## Introduction

Germ cells are essential for fertility, embryonic development, and reproduction. In animals, germ cells undergo a highly dynamic developmental program. Primordial germ cells (PGCs) are first specified in the early embryo and then migrate to the developing gonad. In the gonads, PGCs give rise to germline stem cells that in turn produce oogonia or spermatogonia which later initiate differentiation by the induction of the meiosis program. Sex determination mechanisms in the gonad direct early germ cells to initiate either oogenesis or spermatogenesis, for the generation of functional egg or sperm cells, through intricate mechanisms of differentiation and morphogenesis. Identifying regulators and deciphering the mechanisms by which they control the dynamic multi-step differentiation of germ cells is essential for better understanding of their development, as well as fertility and reproduction.

In zebrafish, the regulation of various steps in germ cell development requires three types of RNA-Protein (RNP) granules. First, RNP granules that contain germ cell fate determinants, termed germ plasm, initially specify PGCs in the embryo, and then maintain their germ fate and are required for their proper migration to the gonad [1]. The germ plasm includes the RNP components Vasa, Dead-end (Dnd), and Bucky ball (Buc) [2,3].A second type of RNP granules is called germ granules and form in oogonia and early differentiating oocytes in the ovary [4]. Germ granules are localized perinuclearly and contain the PIWI interacting RNAs (piRNA) pathway machinery [5], which protects germline nuclei from retrotransposons by processing and cleaving their transcripts [4–8]. Germ granules in most animals [5,7], including zebrafish [9,10], associate with clustered mitochondria and were collectively referred to as nuage. Zebrafish germ granules contain the piRNA proteins Ziwi, Zili, as well as Vasa, and are essential for repression of transposon expression [10].

A third type of RNP granules form in an oocyte membraneless organelle, which is conserved from insects to humans, and called the Balbiani body (Bb) [11–16]. The Bb establishes the oocyte animal-vegetal polarity, by specifying the vegetal pole of the egg and future embryo, which is key for embryonic development [17]. Bb RNP granules include the germ plasm components as well as dorsal determinants [18]. The Bb delivers the germ plasm and dorsal determinants to the oocyte vegetal pole from which they will later act to specify the germline and to establish the embryonic dorsal-ventral axis [18]. In oogenesis, following an early symmetry-breaking event that polarizes Bb granule localization around the centrosome [13], the Bb forms adjacent to the oocyte nucleus and then translocate to the oocyte cortex [19]. At the cortex, the Bb disassembles, docking its RNP granules to the cortex, and thus specifying this region as the oocyte vegetal pole [19]. A fundamental understanding of the regulation of Bb RNP complexes is lacking, and the Bucky ball (Buc) protein is the single known essential Bb protein in any species [13,20]. In *buc* mutants the Bb fails to form, resulting in radially symmetrical eggs and embryonic lethality [20]. Only two other Bb proteins, Tdrd6a and Rbpms2, were shown to affect Bb morphology [21,22]. The Tdrd6a protein interacts with Buc and enhances its granule formation, and the Rbpms2 protein localizes to the Bb, but neither are essential for Bb formation [21,22].

RNP granules are thought to form dynamic molecular condensates, as known for germ plasm and piRNA RNP granules in germ cells and embryos in invertebrates, such as P granules in *C. elegans* and polar granules in Drosophila [23], as well as in somatic mammalian tissues, such as stress granules and P bodies[24]. Accumulating knowledge in invertebrates begin to uncover regulators and decipher mechanisms that control the molecular condensation of granules [25–27], as well as generate the understanding of their hierarchical formation of homotypic and heterotypic RNP complexes [5]. However, our knowledge of the structure, the full repertoire of RNA/protein components, and the regulatory proteins of germ plasm-, piRNA-, and Bb granules is still lacking in vertebrates, including zebrafish.

The Four <u>a</u>nkyrin repeats (ANK), a <u>s</u>terile alpha motif (SAM), and leucine <u>z</u>ipper 1 protein, named Asz1 (previously called GASZ), is a germ cell specific protein that localizes to RNP complexes [28]. In mice gonads, Asz1 is localized to the cytoplasm of pachytene spermatocytes and early spermatids, and is expressed during all stages of oogenesis [28]. In both oocytes and spermatocytes, Asz1 localizes to the perinuclear germ granule RNPs, which contain the machinery of the piRNA pathway [5]. Asz1 co-localizes with the PIWI protein MILI and is required for piRNA processing [29], and it functions similarly in Drosophila [30,31]. piRNA granules associate with clustered mitochondria and are collectively referred to as nuage. Asz1 is important for nuage formation, mitochondrial clustering and transposon repression due to its ability to interact with the mitochondria outer membrane [32,33]. Loss of Asz1 in mice results in increased retrotransposon transcription and male sterility but has no apparent phenotypes in females [29].

In zebrafish, Asz1 was shown to localize in oogonia and early oocytes to germ granules that contain the piRNA proteins Ziwi and Zili, as well as Vasa [13], and in zebrafish and Xenopus, Asz1 localizes to the Bb as well [13,20,34]. Zebrafish Asz1 was shown to undergo concomitant localization dynamics with Bb granules during symmetry-breaking [13], and its localization to the mature Bb was dependent on Buc [20]. Based on the localization of Asz1 to the germ granules and the Bb, and considering its roles in mice [29], we hypothesized that Asz1 could regulate RNP granule dynamics, as well as germ cell and gonad development in zebrafish.

Here, to test this hypothesis, we generated *asz1* mutant fish and determined the roles of Asz1 in germ cell and gonad development. We provide evidence to show that loss of Asz1 exhibit no apparent somatic developmental defects but is essential for germ cell survival and gonad development. *asz1^-/-^* fish developed exclusively as sterile males that completely lacked germ cells. Rare rescue of ovarian development in *asz1;tp53* double mutant fish revealed that Asz1 is similarly essential for oogenesis. Mechanistically, we show that loss of Asz1 results in induction of transposon expression and high levels of germ cell apoptosis, which likely demonstrates a conserved role for Asz1 in the piRNA pathway in zebrafish. Interestingly, germ granules in mutant gonads, exhibited aberrant morphology in which some components appeared to coalesce, while others appeared normal. In contrast, we provide evidence concluding that zygotic Asz1 is not essential in germ plasm granules, and is dispensable for Bb formation. Our work establishes that Asz1 is essential for germ cell and gonad development in both females and males in zebrafish, and reveals differential necessities for this protein in distinct types of RNP granules.

## Results

### Loss-of Asz1 function induces defective germ cell and gonad development

Asz1 is a germline specific piRNA granule protein in the mouse, and a strong candidate regulator of RNP granule biology, as well as germ cell and gonad development in zebrafish. However, the functions of Asz1 in zebrafish have not been addressed. We confirmed by RT-PCR analysis that zebrafish *asz1* is expressed specifically in ovaries and testes, with maternal deposition of transcripts in early embryos (Fig. S1A). To determine the roles of Asz1 in zebrafish, we generated fish carrying a loss-of-function allele of *asz1*.

Consistent with its structure in the mouse, zebrafish Asz1 is a 480 amino acid protein that contains three ankyrin repeats at the N-terminus, a SAM domains in the middle of the protein, and a trans-membrane domain at the C-terminus (Fig. S1D) [34]. We used CRISPR/Cas9 to generate a *loss-of-function* allele, using two independent gRNAs. We targeted exon 6 at the end of the ankyrin repeats (Fig. S1D) with gRNA1, and exon 2 at the middle of ankyrin repeats (Fig. S1D) with gRNA2, and screened for alleles bearing an early termination codon. Despite multiple screening of many F1 families in the F2 and F3 generations, gRNA2 only produced various mismatch mutations that were not predicted to alter the Asz1 protein sequence and function. In contrast, gRNA1 resulted in a four base-pair deletion, generating a premature STOP codon at amino acid 243 (Fig. S1B,D). PCR analysis of *asz1* genomic locus at the region of the gRNA1 sequence, followed by high-resolution gel electrophoresis, confirmed the mutation (Fig. S1C).

This allele is predicted to generate either a truncated protein that lacks the SAM and TM domains and is thus non-functional, or no protein at all in case the transcript undergoes nonsense mediated decay. To distinguish between these two possibilities, we examined *asz1* transcripts by RT-PCR. Below, we determine that loss of *asz1* results in complete loss of germ cells, but that residual germ cells are still present at 4 wpf (Fig. 2-3). Since lack of germ cells precludes their gene expression analysis, we performed this analysis on wt and mutant gonads at 5wpf, and monitored the presence of germ cell by using primers for the germ cell specific marker *vasa*. While *asz1* transcripts were clearly detected in wt and heterozygous gonads, they were almost completely abolished in homozygous *asz1* gonads, despite substantial detection of residual *vasa* expression, which confirms the presence of germ cells in these samples (Fig. S1E). As expected, *asz1* transcripts were also abolished from adult gonads (Fig. S1E). These results suggest that mutant transcripts undergo nonsense mediated decay. Thus, this allele is predicted to be a null *asz1* mutant, and was termed *asz1^huj102^*.

**Figure 1.**
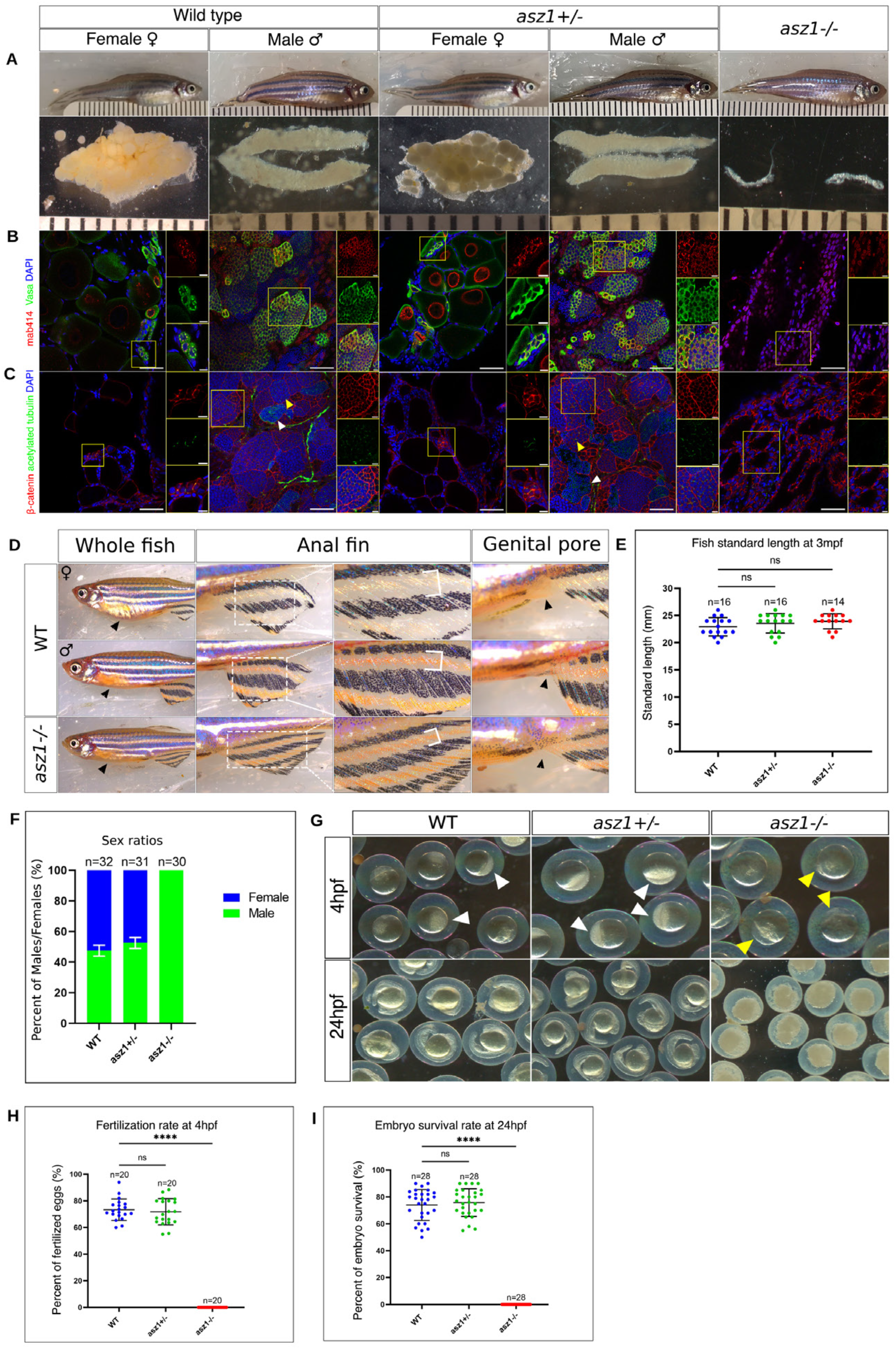
Loss of Asz1 results in germ cell loss and underdeveloped gonads and *asz1^-/-^* fish develop as sterile males. **A.** Representative adult fish at 3 months post-fertilization and their corresponding gonads of the indicated genotypes. Ruler grades are 1mm, showing similar standard lengths (SL) of fish from all genotypes (E), but much smaller gonads of *asz1^-/-^*compared to wt ovaries and testes. **B-C.** Representative confocal images of ovaries and testes of the indicated genotypes as in A, labeled with Vasa (green), mAb414 (red) and DAPI (blue) in B, and with Acetylated tubulin (green), β-Catenin (red) and DAPI (blue) in C. Insets show single and merge channels of magnifications of the yellow boxes in zoomed out images. Scale bars are 50 μm and 10 μm in zoomed out and inset magnification images, respectively. In C, yellow arrowheads indicate zygotene cilia in spermatocytes, and white arrowheads indicate flagella of mature sperm. For each B and C labeling, n= 6 ovaries and 6 testes from wt and *asz1^+/-^*, and 6 underdeveloped gonads in *asz1^-/-^*, from two representative clutches. **D.** Dimorphic external sex criteria in wt, and their phenotypes in *asz1^-/-^* fish. Left panels show larger abdomen in females (arrowheads). Middle panels show the anal fin (right images are zoomed-in magnifications), with more pigmented stripes (white bracket) in the male. Right panels show the larger genital pore in the female (arrowheads). *asz1^-/-^* fish exhibit typical male anatomy of all criteria. The number of analyzed fish is indicated in the plot in F. **E.** A plot of the SL of fish from all genotypes. n=number of fish. SL was not significantly different between genotypes. Bars are mean ± standard deviation (SD). **F.** A plot showing the representative sex ratios in each genotype from two independent clutches as determined by the external sex criteria in D. n=number of fish. Bars are mean ± SD. **G.** Progeny embryos of crosses between wt females and either wt, *asz1^+/-^*, or *asz1^-/-^* males as indicated, at 4 and 24 hpf. At 4 hpf, fertilized embryos exhibit an opaque animal pole which results from cellularization during cleavage stages (white arrowheads), while unfertilized eggs exhibit a transparent acellularized animal pole (yellow arrowheads). At 24 hpf, unfertilized embryos from wt females and *asz1^-/-^* males are lysed (right panel). **H-I.** Dot plots showing the fertilization (H) and survival rates (I) per mating from the crosses in G. n=number of cross pairs. Bars are mean ± SD.

**Figure 2.**
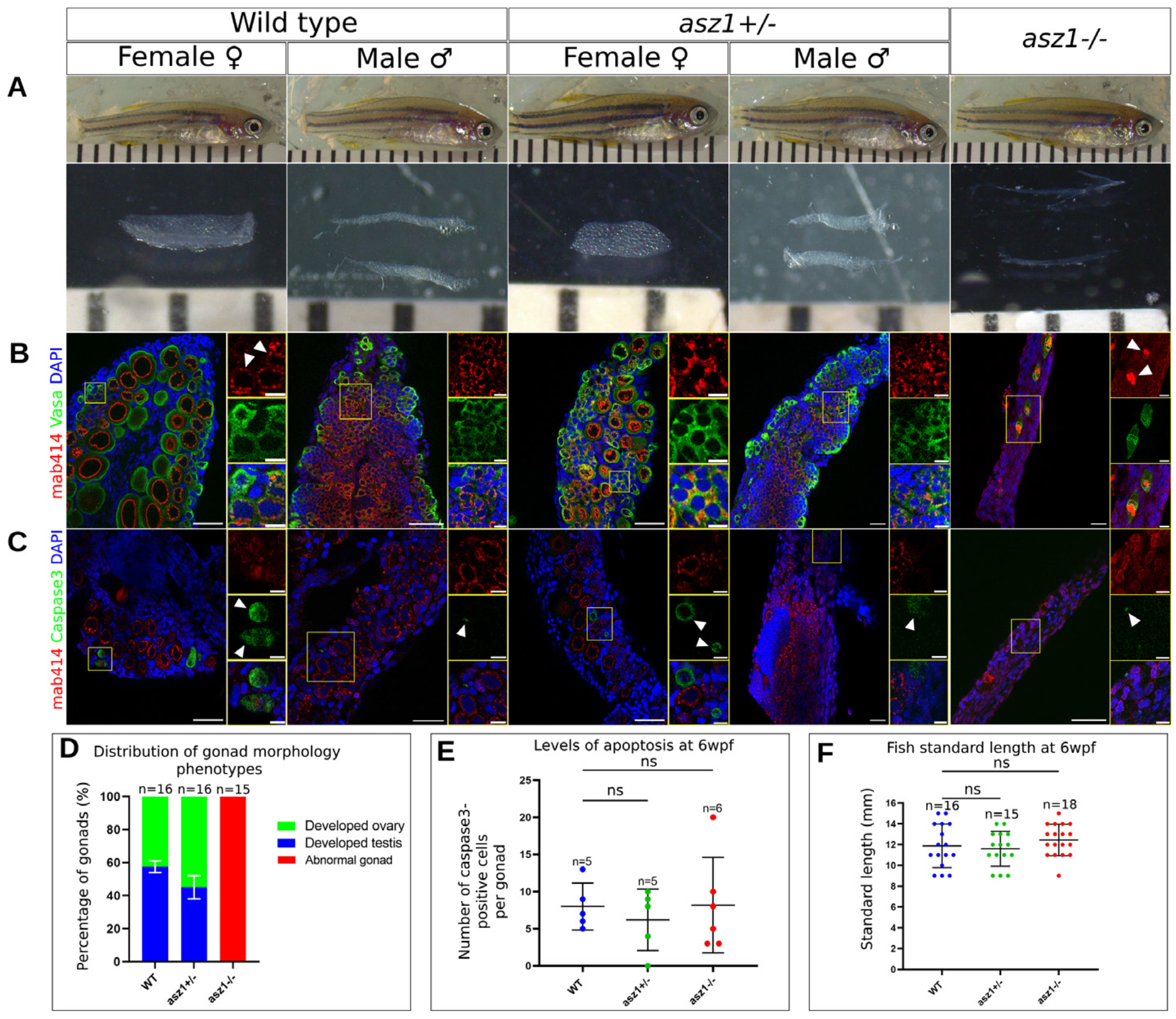
Asz1 is essential for germ cell and gonad development in juvenile post-embryonic stages. **A.** Representative juvenile fish at 6 wpf and their corresponding gonads of the indicated genotypes. Ruler grades are 1mm, showing similar standard lengths (SL) of fish from all genotypes (F), but much smaller, thread-like, gonads of *asz1^-/-^* compared to wt ovaries and testes. The distribution of gonad morphology from two representative clutches is plotted in **D.** n=number of gonads. Bars are mean ± SD. **B.** Representative confocal images of ovaries and testes of the indicated genotypes as in A, labeled with Vasa (green), mAb414 (red) and DAPI (blue). Insets show single and merge channels of magnifications of the yellow boxes in zoomed out images. Scale bars are 50 μm and 10 μm in zoomed out and inset magnification images, respectively. Wt ovaries (n=9) and testes (n=5), as well as *asz1^+/-^* ovaries (n=7) and testes (n=7), exhibited normal developing oocytes and early spermatocytes, as well as oogonia and spermatogonia as indicated by Vasa, with normal perinuclear piRNA granules, as indicated by mAb414 signals (white arrowheads in wt ovary). *asz1^-/-^* gonads (n=13) had only few Vasa-positive germ cells which exhibited abnormal mAb414 signal that appeared as coalesced granules (white arrowheads in *asz1^-/-^* gonads). **C.** Representative confocal images of ovaries and testes of the indicated genotypes as in A, labeled with cCaspase3 (green), mAb414 (red) and DAPI (blue). Insets show single and merge channels of magnifications of the yellow boxes in zoomed out images. Scale bars are 50 μm and 10 μm in zoomed out and inset magnification images, respectively. White arrowheads indicate cCaspas3-positive apoptotic cells. n=5 ovaries and 5 testes (wt), n=5 ovaries and 5 testes (*asz1^+/-^*), n= 6 gonads (*asz1^-/-^*). The number of apoptotic cells per gonad is plotted in **E** and is non-statistically significant between genotypes. n=number gonads. Bars are mean ± SD. **F.** A plot of the SL of fish of all genotypes. n=number of fish. SL was not significantly different between genotypes. Bars are mean ± SD.

**Figure 3.**
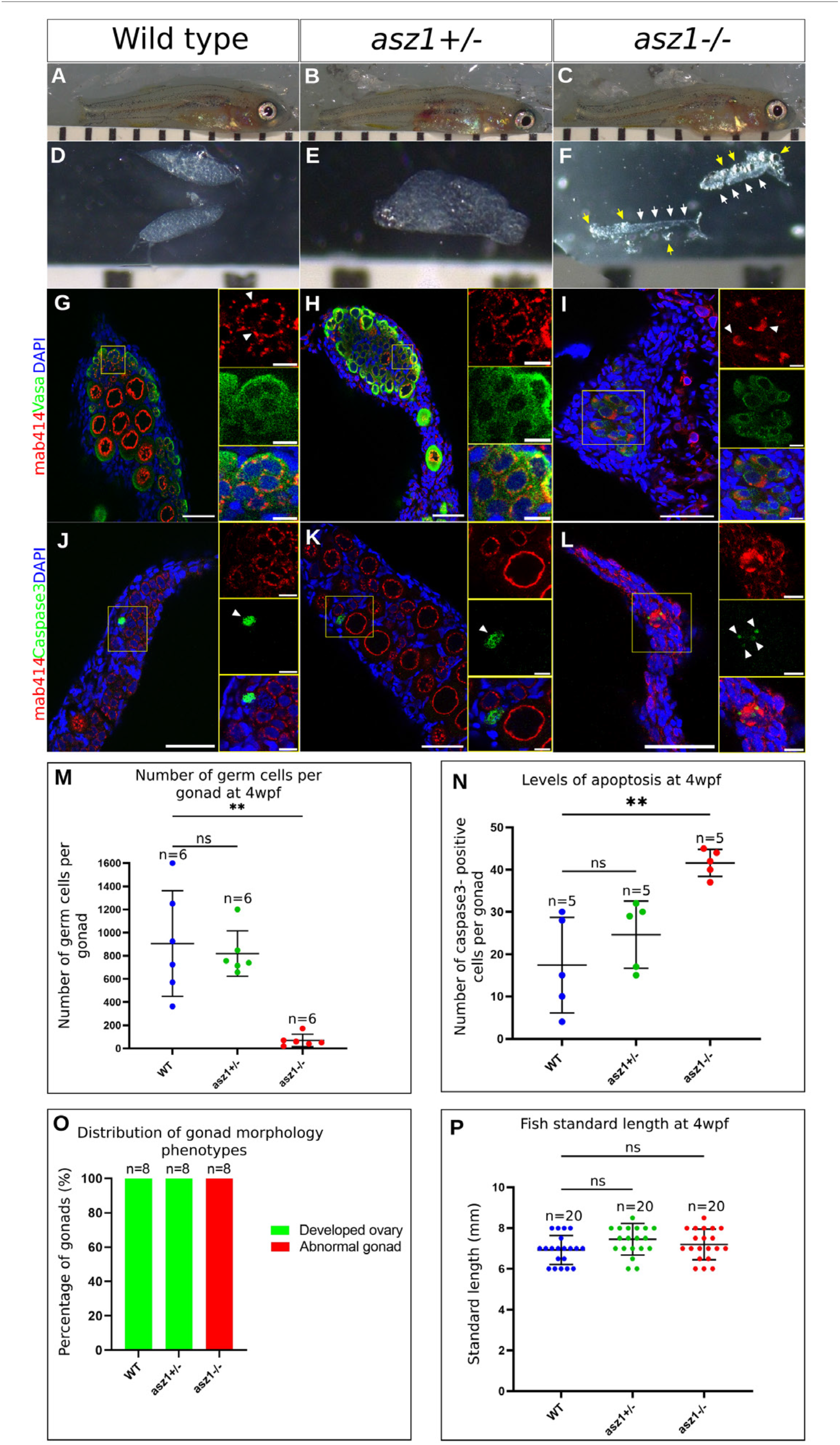
Asz1 is essential for germ cell survival during early gonad development. **A-L.** Fish, gonads and confocal images of labeled gonads are shown as in Fig. 1-2, except fish and their gonads are analyzed at 4 wpf. **D-F.** At 4 hpf, prior to sex determination, all gonads are still developing as ovaries as shown in wt and *asz1^+/-^*. *asz1^-/-^* exhibit much smaller thread like gonads (white arrows in F) already at 4hpf. Yellow arrows in F indicate surrounding bright fat tissue. **G-H.** Developing ovaries in wt and *asz1^+/-^*contain oogonia and early differentiating oocytes (Vasa, green) with normally organized perinuclear granules (mAb414, red), but much fewer germ cells in *asz1^-/-^* that also exhibit coalesced mAb414 signals (white arrowheads in I). n=6 gonads per genotype. **J-L.** cCaspase3 labeling (green) of wt, *asz1^+/-^*, and *asz1^-/-^* gonads. Cells with coalesced mAb414 signals in *asz1^-/-^*gonads (L), which are likely abnormal germ cells, are positive for cCaspase3 (white arrowheads in L). n=5 gonads per genotype. Scale bars in G-L are 50 μm and 10 μm in zoomed out and inset magnification images, respectively. **M.** The number of Vasa-positive germ cells per gonad is plotted for each genotype. n=number of gonads. Bars are mean ± SD. **N.** The number of cCaspase3-positive cells are per gonad is plotted for each genotype. n=number of gonads. Bars are mean ± SD. O. The distribution of normal developing ovaries (as shown in D, E), and abnormal gonads (as shown in F) is plotted per genotype. n=number of gonads. **P.** A plot of the SL of fish from all genotypes. n=number of fish. SL was not significantly different between genotypes, as shown by representative images in A-C. Bars are mean ± SD.

We next tested for phenotypes in *asz1* mutants. Sibling wt, *asz1^+/huj102^*, and *asz1^huj102/huj102^*fish were always raised under identical conditions and fish from all genotypes successfully developed to adulthood similarly. Consistently with the gonad specific expression of *asz1*, *asz1^huj102/huj102^* mutant fish seemed to develop normally with no apparent somatic phenotypes (Fig. 1A). As a more accurate measure, the standard length (SL) in zebrafish is used as a parameter of fish growth in adulthood and during post-embryonic development, and is defined as the distance between the fish snout and the base of its tail [35]. The SL of sibling *wt*, *asz1^+/huj102^*, and *asz1^huj102/huj102^* fish were similar (Fig. 1E), with statistically insignificant variation between sibling genotypes (Fig. 1E).

To test for potential germline loss-of-function phenotypes, we analyzed gonads of *wt*, *asz1^+/huj102^*, and *asz1^huj02/huj102^*juvenile and adult fish (Fig. 1A-C, Fig. S2). We first analyzed the general morphology of the gonads, as well as visualized germ cell and gonad organization by DAPI labeling followed by confocal microscopy (Fig. S2). We analyzed developing gonads at 6 weeks post-fertilization (wpf) (Fig. S2A-J), as well as adult gonads at 3 months post fertilization (mpf) (Fig. 1A-C, Fig. S2K-T).

Wt and heterozygous fish exhibited ovaries and testes with normal morphology as expected. Wt and heterozygous ovaries were at the normal size range in both juvenile and adult stages (Fig. 1A, Fig. S2A, E, K, O). Juvenile ovaries contained developing oocytes (Fig. S2B, F), and adult ovaries contained normal premature opaque (yellowish) st. III oocytes (Fig. 1A, Fig. S2L, P) that were surrounded by follicle cells, as detected by DAPI (Fig. S2L, P). Wt and heterozygous testes exhibited normal size range at both stages (Fig. 1A, Fig. S2C, G, M, Q), as well as developing seminiferous tubules in juveniles (Fig. S2D, H), and fully grown tubules filled with mature sperm in adults, as detected by DAPI (Fig. S2N, R). Adult testes also showed a characteristic milky color, indicative of the presence of sperm (Fig. 1A, Fig. S2M, Q). In sharp contrast, homozygous *asz1^huj102/huj102^* fish exhibited much thinner, smaller and transparent gonads that appeared like threads, with no apparent oocytes or milky color, suggesting the absence of germ cells (Fig. 1A, Fig. S2I,S). DAPI staining confirmed that homozygous gonads did not seem to contain either oocytes or sperm, and appeared as “empty” undeveloped gonads that only contained somatic and stromal cells (Fig. S2J, T). Based on these observations, as well as the loss of *asz1* transcripts, we conclude that the *asz1^huj102^*allele presents a loss-of-function *asz1* mutation (hereafter referred to as *asz1^-^*), which causes defective germ cell and gonad development.

To better understand the consequences of *asz1* loss in gonads, we analyzed adult gonads labeled for additional cellular markers. We co-labeled gonads with the germ cell specific marker Vasa [36,37] and with mAb414. mAb414 detects Phenylalanine-Glycine (FG)-repeat proteins in the nuclear pore complexes on the nuclear envelope, as well as a signal that co-localizes with Asz1, Vasa, and other piRNA machinery proteins in perinuclear piRNA granules in zebrafish gonads [13,38]. Vasa-positive germ cells with normal perinuclear granules (either oocytes or spermatocytes) were abundant in wt and *asz1^+/-^*gonads (insets are zoomed in magnification of yellow boxes) (Fig. 1B). However, Vasa-positive cells were completely absent from *asz1^-/-^* gonads (Fig. 1B), confirming the loss of germ cells in mutants.

We next co-labeled gonads with acetylated tubulin and β-catenin (Fig. 1C). Acetylated tubulin labels the zygotene cilium in zygotene oocytes and spermatocytes [39], marking developing germ cells in meiotic prophase, as well as the flagella in mature sperm (white and yello arrowheads in Fig. 1C). β-catenin marks Adherens junctions on cell membranes and is used to visualize cell boundaries and tissue morphology. Wt and heterozygous testes showed normal cilia (yellow arrowheads) and flagella (white arrowheads) structures in spermatocytes and mature sperm, respectively (Fig. 1C), that were organized normally in clear seminiferous tubules. Additionally, wt and heterozygous ovaries showed normal oocytes at various stages of development, as detected by β-catenin (insets are zoomed in magnification of yellow boxes), including ciliated zygotene oocytes as detected by acetylated tubulin (Fig.1C). In contrast, homozygous mutant gonads, completely lacked either zygotene cilia or flagella, further demonstrating the lack of differentiating oocytes and spermatocytes and the lack of mature sperm (Fig.1C), consistently with the absence of Vasa-positive cells (Fig. 1B). Moreover, β-catenin labeling revealed cellular gaps, that appeared as empty holes in the tissue, which could indicate an earlier loss of germ cells. These results demonstrate striking gonad morphological defects, as well as complete loss of germ cells in adult *asz1* mutant fish, concluding that Asz1 is essential for gonad and germ cell development in zebrafish.

### *asz1* mutants develop as sterile males

The complete lack of germ cells in *asz1* mutants made it challenging to determine the sex of *asz1* mutant fish. In zebrafish mutations that lead to early germ cell loss can result in male sex bias or sex reversal to males [40–42]. Oocytes are required for maintaining the ovary fate in zebrafish by yet unknown mechanisms [43]. This maintenance perishes with oocyte loss and gonads adopt a testis fate [43]. The thin morphology of *asz1* mutant gonads in adult and juvenile stages, resembled testes morphology (Fig. 1A, Fig. S2J, T). Moreover, β-catenin labeled mutant gonads exhibited gaps or holes in the tissue (Fig. 1C) that could represent abnormal seminiferous tubules that lacked germ cells and appeared as empty tubules, as previously shown in *vasa* mutant fish [44], and suggesting that *asz1* mutant gonads develop as defective testes.

To definitively determine the sex of *asz1* mutants, we used three established external sex dimorphic criteria in zebrafish: ***1)*** a larger abdomen in the female (which contains the ovary with large premature and mature oocytes) (Fig. 1D), ***2)*** a dimorphic color of the anal fin which is composed of yellow and black strips in males, and of pale and black stripes in females (Fig. 1D), and ***3)*** the genital pore, which is located at the ventral side of the abdomen anterior to the anal fin and posterior to the anus, and is larger and protrudes more substantially from the body in females compared to males (Fig. 1D) [45]. We thus scored fish sex based on those external criteria. As shown in pooled data from two representative clutches (Fig. 1F), wt (n=32) and *asz1^+/-^* (n=31) fish exhibited approximately 1:1 female-male sex ratios as expected. However, 100% of *asz1^-/-^* fish (n=30) exhibited all three male-characteristic dimorphic criteria, including smaller abdomen (Fig. 1F), more significant yellow color strips (Fig. 1D), and a male-typical genital pore (Fig. 1D). Therefore, based on gonad morphology and external sex criteria, we conclude that loss of Asz1 function results in a complete male bias.

The lack of germ cells in *asz1* mutants predicted that adult males are sterile. To determine fertility, we tested whether *asz1^-/-^* males can fertilize wt eggs and/or produce normally developing embryos. We crossed wt females with either wt, *asz1^+/-^*, or *asz1^-/-^* males (Fig. 1G-I) and determined the fertilization rates at 4 hours post fertilization (hpf) (Fig. 1G-H), as well as embryo survival at 24hpf (Fig. 1G, I). At 4 hpf, wt and *asz1^+/-^* males crossed to wt females resulted in normal 75%±5% and 73%±7 fertilization rate, respectively, as evident by the more opaque color of the animal pole that results from cellularization during cleavage stages, as opposed to the transparent single animal cell in non-fertilized eggs (n=20 pairs of wt female and wt male, n=20 pairs of wt female and *asz1^+/-^* male) (Fig.1G-H). In contrast, in crosses of wt females with *asz1^-/-^* males (n=20 pairs), normal eggs were laid, but none (0%) were fertilized (Fig. 1G-H). At 24 hpf, embryo survival rates were 73%±10 from wt males, 75%±12 from *asz1^+/-^* males, and 0% from *asz1^-/-^* males (n=28 pairs of each genotype combinations) (Fig. 1I). These results concluded that *asz1* mutant males are sterile as expected from their lack of sperm. Altogether our data establishes that the Asz1 protein is essential for germ cell and gonad development, and that its loss is detrimental to fertility and reproduction.

### Asz1 is essential for early gonad and germ cell development in the juvenile fish

To understand the developmental defects in *asz1* mutant gonads, we sought to determine the timeframe during gonad and germ cell development at which Asz1 functions are essential. We thus backtracked *asz1* phenotypes to juvenile stages and analyzed developing gonads at 6wpf. Zebrafish sex determination occurs in juvenile stages, and at 6wpf, both developing ovaries and testes can be captured, as was the case in wt and *asz1^+/-^* fish (Fig. 2A). However, we found that the developmental defects in *asz1* mutants were already detected at these stages, and *asz1^-/-^* gonads appeared as extremely smaller, thinner, and underdeveloped gonads compared to wt (Fig. 2A). Notably, the SL of all sibling genotypes were similar with no statistically significant differences (Fig. 2F), further ruling out indirect growth defects during zebrafish development.

To monitor germ cell development, we labeled gonads with Vasa and mab414. Vasa labeling detected numerous germ cells at various developmental stages in wt and heterozygous testes (n=5 and 7 testes, respectively) (Fig. 2B), as well as in wt and heterozygous ovaries (n=9 and 7 ovaries, respectively) (Fig. 2B). Strikingly, in *asz1* mutant gonads, only few germ cells were detected (n=13 gonads) (Fig. 2B). Furthermore, in contrast with the detection of mAb414 in normally organized perinuclear granules in wt and *asz1^+/-^* gonads (Fig. 2B-C, white arrowheads in B) [13], in the few existing germ cells in *asz1^-/-^* gonads, mAb414 signal was detected as an abnormal and polarized large aggregate in the cell (Fig. 2B-C, white arrowheads in B). The mis-organized mAb414 signal could indicate a piRNA granule defect in *asz1* mutants. However, Vasa-positive granules remained normally perinuclear (Fig. 2B).

The plot in Figure 2D summarize these phenotypes. Based on the Vasa and mAb414 analyses above, we categorized all analyzed gonads as “normal testes” or “normal ovaries”, if they showed normal number and developmental stages and organization of spermatocytes or oocytes, respectively (as shown in wt and *asz1^+/-^* in Fig 2A-B). We defined thinner gonads with no or very few germ cells, as “abnormal gonads”, which considering the male bias in *asz1* mutants likely represent underdeveloped testes (as shown in *asz1^-/-^* in Fig. 2A-B). While wt (n=16) and *asz1^+/-^* (n=16) gonads developed as either normal ovaries or normal testes (Fig. 2D), with no detected abnormal gonads, *asz1^-/-^* gonads (n=15) were all classified as “abnormal gonads” and did not include any normal ovaries or testes (Fig. 2D). These results conclude that Asz1 is essential for early germ cell development already in the juvenile gonad.

The striking and early loss of germ cells in 6 wpf *asz1^-/-^*gonads suggested that germ cells could be lost by apoptosis. To monitor apoptosis, we labeled gonads from all three genotypes with the apoptosis marker cleaved Caspase3 (cCaspas3) (Fig. 2C, white arrowheads). We detected a similar number of cCaspase-positive apoptotic cells in wt (8±3 apoptotic cells per gonad, n=5 gonads), *asz1^+/-^* (7±4 apoptotic cells per gonad, n=5 gonads), and *asz1^-/-^* (8±7 apoptotic cells per gonad, n=6 gonads) gonads, with statistically insignificant differences (Fig. 2E). This suggest that germ cells are either lost by another mechanism, or that they have already been lost by apoptosis earlier in gonad development. We therefore next analyzed gonads at earlier developmental stages.

### Asz1 is essential for germ cell survival at the onset of gonad development

To further backtrack the loss of germ cells and the developmental defects in the *asz1* mutants, we analyzed gonads closer to their developmental onset, at 4wpf (Fig. 3). Consistently with our previous results at later stages, the SL of fish from all genotypes was similar (Fig. 3P), thus concluding that this mutation does not seem to affect zebrafish somatic development (Fig. 3A-C, P). Prior to sex determination in zebrafish, gonad develop as ovaries, and upon sex determination they either continue to develop as ovaries, or convert to testis development [44] . At 4wpf, preceding sex determination, all gonads still develop as early ovaries [46]. Indeed, wt and *asz1^+/-^* gonads all appeared as normal early developing ovaries (Fig. 3D, E). However, *asz1^-/-^* gonads were again much smaller and thinner compared to siblings (Fig. 3F), demonstrating that Asz1 functions are required from the onset of gonad development.

To examine the cellular phenotypes at these stages, we labeled gonads with Vasa and mAb414. To test for potential germ cell loss at 4 wpf, we determined the number of germ cells per gonad in each genotype, by counting Vasa-positive cells. Wt (n=6) and *asz1^+/-^*(n=6) gonads contained 920±470 and 820±220 germ cells per gonad, respectively (Fig. 3M). In sharp contrast, *asz1^-/-^* gonads (n=6) contained only 90±90 germ cells per gonad (Fig. 3M), further confirming the early loss of germ cells in this mutant. Interestingly, in existing germ cells in *asz1^-/-^* gonads, mAb414 labeling exhibited the same abnormal polarized aggregation phenotype as in 6 wpf gonads. Such aggregates were not detected in wt and *asz1^+/-^* gonads, which showed normal perinuclear granules (Fig. 3G-I).

We next examined apoptosis in 4wpf gonads, as labeled by cCaspase3. We counted cCaspas3-positive cells per gonad in each genotype, and found that apoptosis levels were abnormally and statistically significantly higher in *asz1^-/-^* gonads. *asz1^-/-^* mutant gonads (n=5) contained as high as 42±3 apoptotic cells per gonad, compared to only 17±11 and 25± 7 apoptotic cells per gonad in wt (n=5), and *asz1^+/-^* gonads (n=5), respectively (Fig. 3N). These results conclude that upon loss of Asz1 functions, germ cells are cleared by apoptosis early in gonads development, and that Asz1 is essential for their survival. However, the number of apoptotic cells detected in mutant gonads at theses stages (∼42) cannot account for the presumptive loss of hundreds of germ cells as shown by our counting of Vasa-positive germ cells above (Fig. 3M; compare the ∼920 cells/gonad in wt to ∼90 cells/gonad in *asz1^-/-^* mutants). It is possible that germ cells are lost by other mechanisms in addition to apoptosis in *asz1* mutants. Alternatively, but non-mutually exclusively, these observations suggest that clearance by apoptosis is continuous over the period of gonadogenesis in *asz1* mutants, and that our analyses captured snapshots of apoptosis events at specific timepoints. Supporting this notion, in *asz1* mutant gonads, the majority of germ cells that were detected with cCaspase3 labeling (as also identified by the aggregated mAb414 signal) were detected as fragmented signals, that may represent either early or late time points during the apoptosis process of these specific cells. These results conclude that Asz1 is essential for germ cell survival and gonad morphogenesis already at the onset of gonad development.

### Zygotic Asz1 is dispensable for PGC specification and migration to the gonad, but essential for germline transposon silencing in the gonad

We next investigated the potential mechanism that results in germ cell loss in *asz1* mutant gonads. The essential Asz1 functions in germ cell survival as early as 4 wpf, suggest that Asz1 functions could be required in the developing gonad, or during earlier stages. Prior to germ cell differentiation in the early developing gonad, germ cells and somatic cells of the gonad develop from different embryonic origins. Germ cell precursors, called primordial germ cells (PGCs), are specified by maternal inheritance of germ plasm during early embryonic development, and the gonad develops at later stages at the urogenital ridge [47]. PGCs migrate to the future developing gonad and populate it [47]. In zebrafish, PGCs migrate to the region of the future developing gonad by 24 hours post-fertilization and populate it by 7 days post-fertilization [1,15]. It is thus possible that the lack of germ cells observed in early developing *asz1* gonads represent an earlier loss of PGCs, including their aberrant specification, and/or failure to migrate to the gonads. We thus wanted to determine whether Asz1 is required for PGCs specification or migration, prior to gonad development.

To address potential earlier phenotypes, we labeled PGCs using the germline specific marker Vasa, in wt, *asz1^+/-^*, and *asz1^-/-^* embryos at 24 hpf (Fig. 4A-B), as well as in larvae at 7 dpf (Fig. 4C-D). We counted Vasa-positive PGCs per embryo at 24 hpf (Fig. 4A; lateral view is shown), detecting 23±6, 19±6, and 22±6 PGCs per embryo in wt (n= 22 embryos), *asz1^+/-^* (n= 31 embryos), and *asz1^-/-^* (n=9 embryos) embryos, respectively, with no statistically significant differences between the genotypes (Fig. 4B). The number of PGCs per larvae at 7dpf was consistently similar between genotypes with no statistically significant differences (Fig. 4D). We detected 21±3, 21±4, and 21±2 PGCs per larvae in wt (n=17), *asz1^+/-^* (n=17), and *asz1^-/-^* (n=14) larvae, respectively (Fig. 4C-D). Thus, in zygotic *asz1* mutants, PGCs seem to be specified at normal numbers [48] and to properly migrate to the gonad, demonstrating that zygotic Asz1 is not a major functional regulator of PGC specification and migration in zebrafish. These experiments conclude that in *asz1* mutants, germ cells are lost directly in the gonad in a developmental timeframe between 7 dpf and 4 wpf.

**Figure 4.**
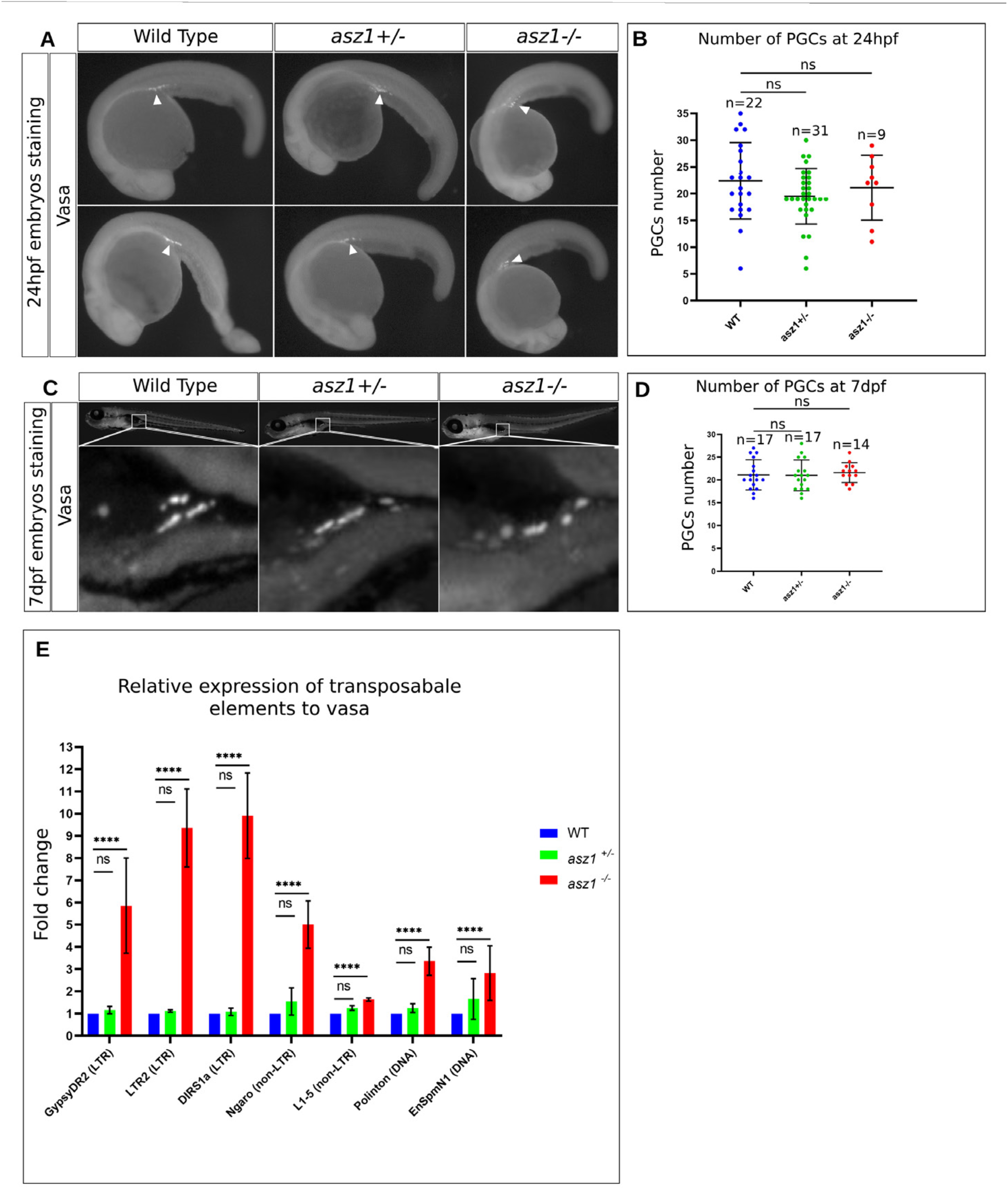
Zygotic Asz1 is not required for PGC specification and migration, but is required for suppression of transposon expression in gonadal germ cells. **A.** Vasa labeling in 24 hpf wt, *asz1^+/-^*, and *asz1^-/-^* embryos shows normal localization of PGCs to the presumptive gonad region (arrowheads). **B.** The number of PGCs per embryo as shown in A is plotted for each genotype and shows non-significant variation across genotypes. n=number of embryos. Bars are mean ± SD. **C.** Vasa labeling in 7 dpf wt, *asz1^+/-^*, and *asz1^-/-^* larvae shows normal localization of PGCs to the forming gonad region. Bottom images are zoom-in magnifications of the boxes in larvae in the top panels. **D.** The number of PGCs per larva as shown in C is plotted for each genotype, is consistent with the number of PGCs detected at 24 hpf in A-B, and shows non-significant variation across genotypes. n=number of embryos. Bars are mean ± SD. **E.** RT-qPCR analysis of the expression of germline transposons in 5 wpf gonads from each genotype (n=50 gonads per genotype), as normalized for RNA loading by *β-act* as well as for germ cells by *vasa* expression (see methods). Bars are mean ± SD from biological duplicates, each made with technical replicates. Statistical analyses were tested by ANOVA.

Having determined that Asz1 is essential directly in the developing gonad and considering the known roles of Asz1 in piRNA processing in Drosophila and mice [29–31], we hypothesized that Asz1 could function similarly in zebrafish. We reasoned that loss of *asz1* would result in de-repression of retrotransposon expression, and that consequently, over-expression of transposons in *asz1* gonads could account for the severe loss of germ cells we observed. Several germline specific transposon in zebrafish, including the long terminal repeat (LTR) transposons *LTR2* and *GypsyDR2*, the non-LTR transposons *DIRS1a*, *Ngaro*, and *L1-5*, and the DNA transposons *Polinton* and *EnSpmnN1*, were shown to be upregulated in *ziwi* and *zili* mutant gonads [4,10]. We therefore monitored the expression of those germline specific transposons in wt, *asz1^+/-^*, and *asz1^-/-^* gonads.

To monitor transposon expression, we aimed to analyze gonads at early stages that precede the complete loss of germ cells in *asz1^-/-^* gonads. Adult mutant gonads completely lacked germ cells (Fig. 1B), and germ cells were scarce starting as early as juvenile stages at 6 wpf (Fig. 2B), which would preclude analyses of their transposon expression. We therefore analyzed gonads at 5 wpf, when *asz1* phenotypes were clearly detected but gonads still contained germ cells. We extracted total RNA from wt, *asz1^+/-^* and *asz1^-/-^* gonads and analyzed the expression levels of the above germline specific transposons by RT-qPCR. Because of the small size of gonads at 5 wpf, and in particular of the underdeveloped *asz1^-/-^* gonads, we extracted RNA from 50 gonads per genotype to obtain sufficient RNA yield.

Normalized expression of all transposons examined and their relative expression to the expression of the germ cell specific marker *vasa* showed background levels in wt gonads, which were unaltered in *asz1^+/-^* gonads (Fig. 4E). In sharp contrast, *asz1^-/-^* gonads exhibited higher expression levels of all transposons examined, ranging from mild upregulation in the case of *L1-5*, to ∼3-10-fold higher expression of *GypsyDR2*, *DIRS1a*, *Ngaro, Polinton,* and *EnSpmnN1* (Fig. 4E). These results demonstrate that upon *asz1* loss of function, retrotransposons are derepressed and over-expressed, concluding that Asz1 is required for germline transposon silencing, likely through the piRNA pathway, consistently with its functions in Drosophila and mice [29–31]. They further propose that the loss of germ cells in *asz1^-/-^* gonads by apoptosis, is likely due to miss-regulated transposon activity in germ cells, as is the case in *ziwi* loss of function mutant gonads in zebrafish [10]. Based on these data, we conclude that Asz1 is essential for early germ cell survival, as well as for spermatogenesis in zebrafish, and likely functions in the piRNA pathway, as in Drosophila and mice.

### Partial rescue of ovarian development in *asz1;tp53* double mutant fish reveals that Asz1 is essential for oogenesis

In zebrafish oocytes, Asz1 localizes to perinuclear piRNA granules together with the major piRNA enzymes Zili and Ziwi [13]. Asz1 also undergoes polarization dynamics with Bb granules during oocyte symmetry breaking [13], and later localizes to the mature Bb [13]. These observations suggest that Asz1 could function in oogenesis. However, in *asz1* mutants, the severe and early loss of germ cells, together with gonad development as underdeveloped testes, precluded analyses in ovaries. To circumvent this issue, we attempted to rescue ovarian development by crossing the *asz1* mutant to *tp53* mutant fish. In zebrafish, the conversion of ovaries to testes during sex determination requires oocyte apoptosis [44]. In several mutants with severe oocyte defects that induce oocyte apoptosis, rescue from apoptosis on a *tp53* mutant background rescued female and ovarian development [40], enabling functional studies in oogenesis. However, this is not always the case, and in some mutant lines loss of *tp53* failed to rescue ovarian development [49] .

We generated *asz1^+/-^;tp53^+/-^* double mutant fish and attempted to rescue ovarian development by in-crosses to generate *asz1^-/-^;tp53^-/-^* progeny for analyses at juvenile stages. We have performed 17 rounds of in-crosses between 40 individual double heterozygous fish. Progeny juveniles carried the two mutations in nine combinations of genotypes that segregated in the expected Mendelian ratios. We examined whether gonads in *asz1^-/-^;tp53^-/-^*progeny formed developing ovaries or testes, based on gonad morphology, as well as Vasa and mAb414 labeling (Fig. S3C-H). Despite tremendous effort, at 5 wpf from 15 rounds of in-crosses, gonads of all *asz1^-/-^;tp53^-/-^*fish (n=180 gonads) exhibited the *asz1^-/-^* mutant phenotype, and appeared like underdeveloped testes (Fig. S3I). However, in two in-cross rounds we could recover 6 gonads (4 and 2 per in-cross) that appeared as ovaries (Fig. S3A-B; 2 ovaries from the cross in A, and 4 from the cross in B).

In the crosses that yielded juvenile progeny with rescued ovaries, wt, *tp53^+/-^, tp53^-/-^, asz1^+/-^*, *asz1^+/-^;tp53^+/-^,* and *asz1^+/-^;tp53^-/-^* exhibited normal ratios of ovaries and testes (Fig. S3A-B). *asz1^-/-^*, and *asz1^-/-^;tp53^+/-^* exhibited defective testes as shown thus far for *asz1* single mutants (Fig. S3A-B). However, in *asz1^-/-^;tp53^-/-^*, we detected three different gonadal morphologies of normal and defective juvenile ovaries and testes (Fig. S3C-H), as we previously described [39]. 60% of gonads (n=10) exhibited the normal *asz1* defective testes morphology, where testes are thin and contain only a few germ cells, and we termed this category “defective testes” (Fig. S3G-H). 20% of gonads appeared similar to the defective testes, but contained oogonia and no further progressing stages of oogenesis, which we termed “underdeveloped ovaries” (Fig. S3E-F). The remaining 20% of gonads contained progressing oocytes and were termed “developing ovaries” (Fig. S3A-D). The total of 6 rescued ovaries from both crosses described above, belong to this category. The SL of these fish at 5wpf was consistent between all nine genotypes (Fig. 5O). Thus, loss of *tp53* only rarely and partially rescued female development in *asz1* mutant fish.

**Figure 5.**
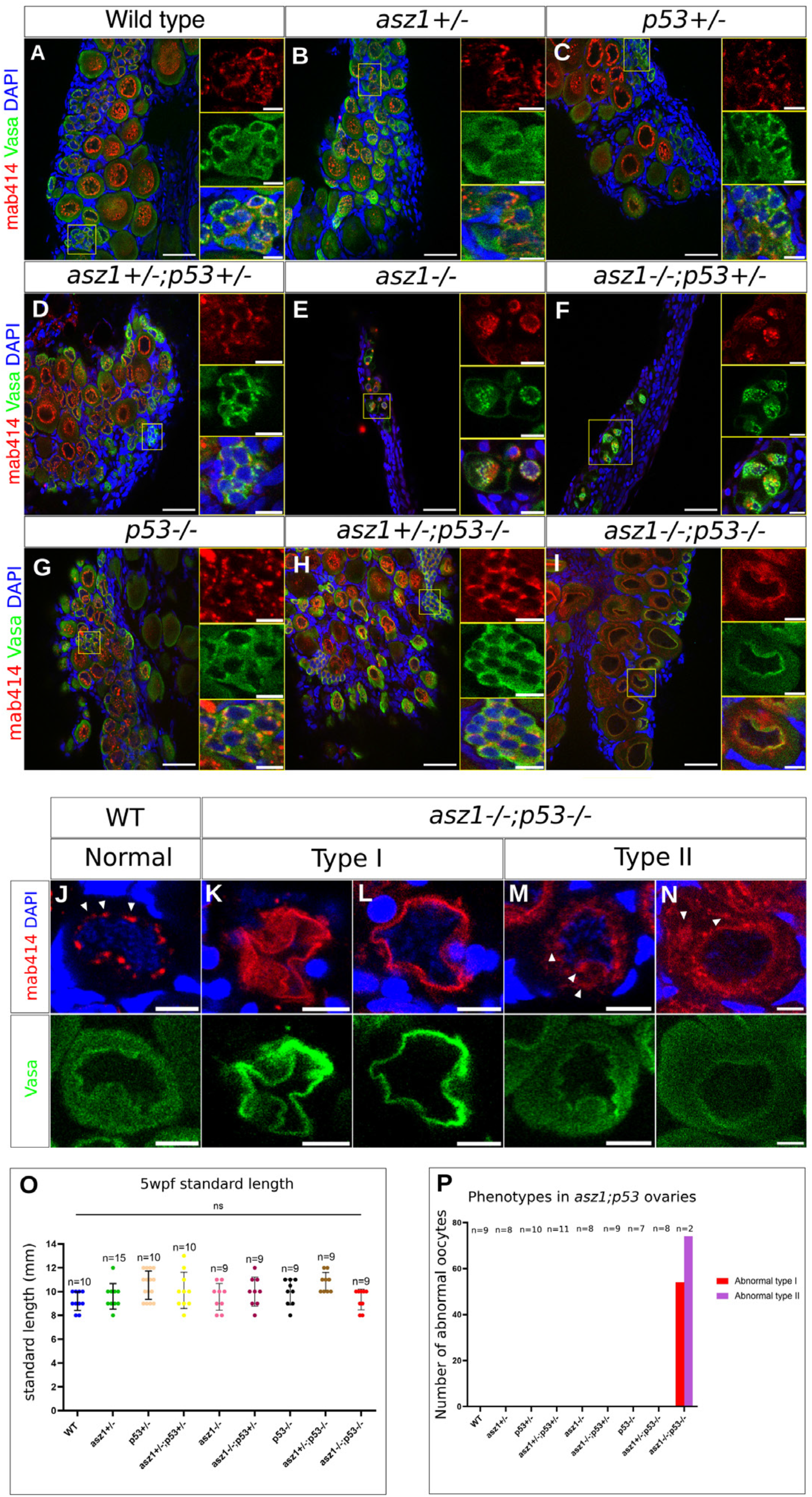
Asz1 loss of function partially rescued ovaries. **A-I.** Ovaries of the indicated genotypes were labeled with Vasa (green), mAb414 (red), and DAPI (blue). Images show representative general morphology of gonads. Right panels are single and merged channels zoomed-in images of the yellow boxed regions in the left panels. Scale bars in G-L are 50 μm and 10 μm in zoomed out and inset magnification images, respectively. Ovaries of all genotypes exhibit normally developing oocytes and ovarian morphology, except the following. asz1^-/-^ and *asz1^-/-^; tp53^+/-^* gonads exhibited the defective testes morphology as shown in previous figures. *asz1^-/-^;tp53^-/-^*ovaries showed defective oocytes, with abnormal nuclear morphology mAb414 granule signals, as detailed in J-N below. n=7-11 gonads per genotype, and 2 ovaries in *asz1^-/-^;tp53^-/-^*, see P below. **J-N.** Images of normal oocytes in the wt (J), and representative defective oocytes in *asz1^-/-^;tp53^-/-^*ovaries (K-N) from panels A, I, showing mAb414 and DAPI in the top panels and Vasa in the bottom panels per oocyte. Two representative images of defective type I oocytes that exhibit abnormal, seemingly collapsed, morphology are shown in K-L. Two representative images of defective type II oocytes with ectopic mAb414 signals that appear as coalesced mis-organized cytoplasmic aggregates are shown in M-N (arrowheads), as opposed to perinuclear granules in wt (arrowheads in J). The distribution of these phenotypes in gonads is plotted in P. Scale bars are 10 μm. **O.** A plot of the SL of fish from all genotypes. n=number of fish. SL was not significantly different between genotypes. Bars are mean ± SD. **P.** The number of defective type I (red) and type II (purple) per gonad is plotted for each genotype. Both phenotypes were only detected in *asz1^-/-^;tp53^-/-^*ovaries. n=number of gonads.

Nevertheless, we analyzed the partially rescued *asz1^-/-^;tp53^-/-^* gonads to obtain insight on potential Asz1 functions in oogenesis. Interestingly, *asz1^-/-^;tp53^-/-^* developing ovaries contained oocytes with abnormal morphology, as detected by Vasa and mAb414 labeling (Fig. 5). Wt, *tp53^+/-^, tp53^-/-^, asz1^+/-^*, *asz1^+/-^;tp53^+/-^,* and *asz1^+/-^;tp53^-/-^* ovaries were normal and contained Vasa-positive oogonia and progressing oocytes with appropriate morphology (Fig. 5A-D, G-H). Oogonia and progressing oocytes exhibited mAb414-positive piRNA granules that were normally organized perinuclearly (Fig. A-D, G-H). *asz1^-/-^* and *asz1^-/-^;tp53^+/-^* gonads were all defective testes, with few germ cells and abnormal mAb414-positive piRNA granule distribution (Fig. 5E-F), as shown for *asz1^-/-^* thus far. However, *asz1^-/-^;tp53^-/-^*ovaries contained progressing of two defective types (n=2 ovaries). Type I defective oocytes exhibited extremely abnormal morphology, where the cytoplasm and nucleus appeared collapsed (Fig. 5K-L, P). Type II defective oocytes in double mutant ovaries exhibited mAb414-positive granules that were either ectopically distributed in the cytoplasm or coalesced into a large aggregate (Fig. 5M-N, P, white arrowheads). We never detected neither phenotype in normal ovaries in the wt, or all other genotypes (Fig. 5P), demonstrating clear defects specifically in *asz1^-/-^;tp53^-/-^*ovaries.

Abnormal oocytes in *asz1^-/-^;tp53^-/-^* developing ovaries likely represent defective oocytes that were rescued from apoptosis, but fail to normally progress through oogenesis due to the loss of Asz1 functions. It is very likely that Asz1 functions in the piRNA pathway in oocytes as well (see discussion). While despite many efforts, we could only recover partial rescue with low number of *asz1^-/-^;tp53^-/-^* ovaries, which preclude further analyses, the above phenotypes conclude that similar to its requirement in spermatogenesis, Asz1 is also essential for oogenesis.

### Asz1 is not essential for Balbiani body formation

Since Asz1 localizes to the Bb in both fish and frogs [13,28], and exhibits co-polarization with Bb granules in zebrafish [13], we wanted to examine whether Asz1 is required for Bb formation. To address this in *asz1* loss of function ovaries, we analyzed Bb formation in all wt, *asz1* and *asz1;tp53* genotype combinations, including four of the *asz1^-/-^;tp53^-/-^*ovaries from the above rescue attempts. To obtain a representative view of Bb RNP granules, we examined the Buc protein, which localizes to the Bb and is essential for its formation [20], as well as the *dazl* Bb mRNA [13]. Buc and *dazl* localize to the Bb during its early formation in the nuclear cleft [20] and through the mature Bb [13,20].

We found that both Buc (n=2 ovaries) and *dazl* (n=2 ovaries) localized normally to the forming Bb in *asz1^-/-^;tp53^-/-^* ovaries, similar to wt (Fig. 6A-D, Fig. S4A, I, Fig. S5A, I), as well as all other genotypes (Fig. S4B-D, G-H, Fig. S5B-D, G-H; n=3-6 gonad per genotype for Buc and 3-5 gonads per genotype for *dazl*), except for *asz1^-/-^*, and *asz1^-/-^;tp53^+/-^*. These latter genotypes developed as defective testes as described thus far (Fig S4E-F, Fig S5E-F) and do not form the Bb, which is a female oocyte-specific organelle. These experiments demonstrate that Asz1 is not essential for the formation of the Bb. Since Asz1 localizes to the Bb, it is still possible that it is required for Bb RNP structure or functions that do not affect Buc and/or *dazl* localization to the Bb, or that it serves redundant functions. Interestingly, together with the likely roles of Asz1 in piRNA granules that we showed above (Fig. 2-4), these experiments suggest differential necessities for Asz1 in distinct RNP granules in the germline.

**Figure 6.**
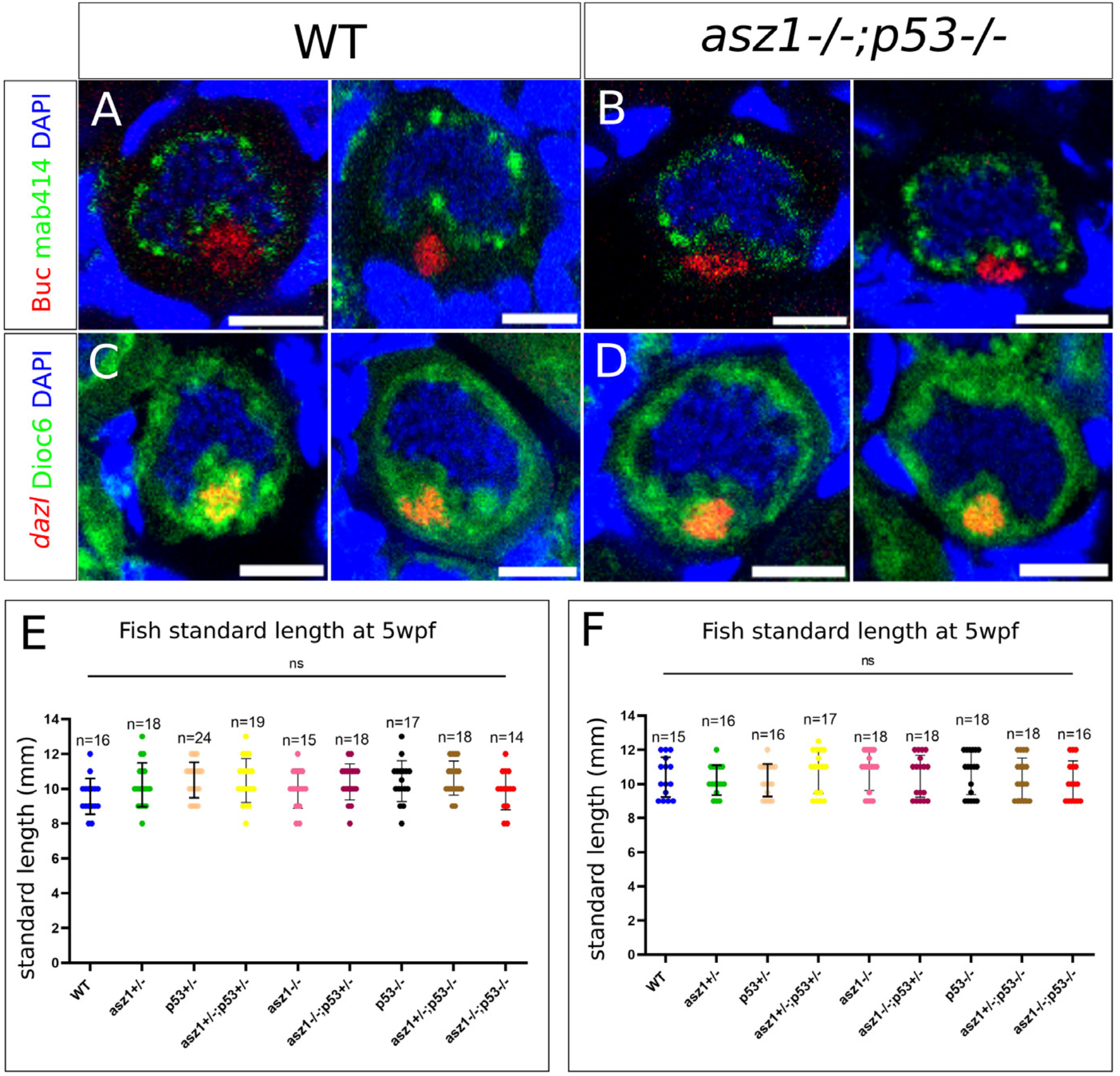
Asz1 is not required for Bb formation. **A-B.** The Buc protein (red) shows normal localization in the forming Bb in the nuclear cleft (mAb14, green; DAPI, blue). n=5 wt ovaries and 2 *asz1^-/-^;tp53^-/-^*ovaries. Representative images of all gonads are shown in Fig. S4. **C-D.** The *dazl* mRNA (HCR-FISH, red) shows normal localization in the forming Bb in the nuclear cleft (cytoplasm labeled with DiOC6, green; DAPI, blue). n=3 wt ovaries and 2 *asz1^-/-^;tp53^-/-^*ovaries. Representative images of all gonads are shown in Fig. S5. Scale bars in A-D are 10 μm. **E-F.** Plots of the SL of fish from all genotypes for the Buc experiment (E) and the *dazl* experiment (F). n=number of fish. SL was not significantly different between genotypes. Bars are mean ± SD.

## Discussion

In this work, we uncover the Asz1 protein as an essential regulator of germ cell and gonad development in zebrafish (Fig. 7). We show that zygotic loss of *asz1* results in severe loss of germ cells in developing gonads between 7dpf and 4 wpf. *asz1^-/-^* fish develop exclusively as sterile males with underdeveloped testes that contain no germ cells (Fig. 7A), concluding that Asz1 is further essential for spermatogenesis. Partial rescue of ovary development revealed that Asz1 is essential for oocyte development as well.

**Figure 7.**
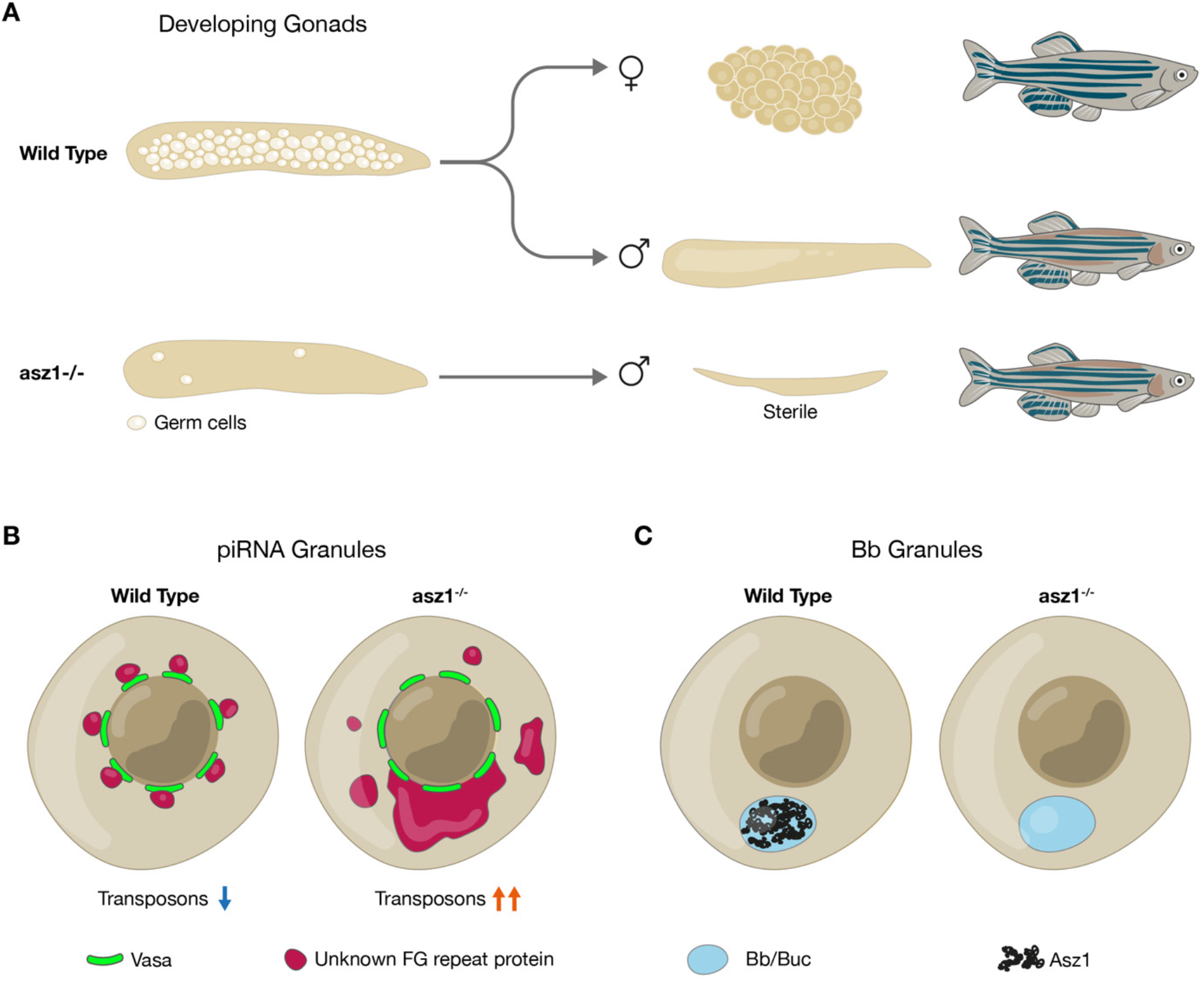
Schematics of the germline developmental requirement for Asz1 functions. **A.** Wt juvenile gonads contain developing germ cells and give rise to ovaries and testes and fertile adult fish. Upon loss of Asz1, germ cells in juvenile gonads are lost at least partly by apoptosis, resulting in underdeveloped testes-like gonads that completely lack germ cells in the adult, and leading to development fish as sterile males. **B.** Asz1 is essential for suppression of transposon expression very likely by acting in the piRNA pathway as shown in Drosophila and mice (blue and orange arrows). Loss of Asz1 also results in mis-organization of piRNA granules as detected by a yet unknown FG-repeat protein (maroon) of these granules. **C.** In contrast with piRNA granules, Asz1 (black) localizes to the Bb (light blue), but upon its loss, Bb granules are normally intact. These observations reveal differential necessities for Asz1 in distinct types of germline granules in zebrafish.

We provide evidence to demonstrate that Asz1 is essential for germ cell survival during early gonad development, at least partly by protecting germ cells form miss-regulated transposon activity and apoptosis, and very likely through the piRNA pathway (Fig. 7B). In early developing gonads, we detected a 2 to 10-fold increase in germ cell expression of LTR-, non-LTR- and DNA transposons (Fig. 4E), concomitantly with germ cell clearance by apoptosis (Fig. 3N), strongly arguing for a role for Asz1 in the piRNA pathway. Asz1 function in the piRNA pathway in zebrafish is consistent with its known roles in piRNA processing in Drosophila and mice [29–31]. The development of *asz1* mutants as males is very likely due to the early loss of germ cells, since in zebrafish oocytes are required for maintaining the female fate, and upon their loss fish develop as males [43]. This is consistent with the male phenotypes in other piRNA mutants in zebrafish, where germ cells are lost by apoptosis at pre-meiotic stages [10], or at early [4], or later [6] meiotic differentiation due to miss-regulation of transposon expression. Notably, our data cannot exclude that germ cells are lost by other mechanisms in addition to apoptosis in *asz1* mutants, and future investigation is needed to reveal additional roles for Asz1 in zebrafish germ cells. Nevertheless, overall, our data establish that *asz1^-/-^* germ cells fail to undergo spermatogenesis.

We further provide evidence to demonstrate that Asz1 is essential for oogenesis. We generated *asz1^+/-^;tp53^+/-^* fish and performed demanding attempts to rescue ovarian development and obtain *asz1* loss-of-function ovaries. We performed 17 rounds of in-crosses between 40 individual double heterozygous fish and out of a total of 200 double homozygous *asz1^-/-^;tp53^-/-^* gonads of progeny at 5wpf, we could only recover 6 developing ovaries. Those 6 ovaries exhibited abnormal oocytes, demonstrating the need for Asz1 for proper oogenesis. Abnormal oocytes in *asz1^-/-^;tp53^-/-^* developing ovaries likely represent defective oocytes that were rescued from apoptosis, but fail to normally progress through oogenesis due to the loss of Asz1 functions.

While the lack of a sufficient number of rescued ovaries prevented direct testing of transposon expression by RT-qPCR, it is very likely that Asz1 functions in the piRNA pathway in oocytes as well. The germline specific transposons above were shown to be expressed similarly in both testes and ovaries in zebrafish [4]. Furthermore, silencing of germline transposons by the piRNA pathway is essential in zebrafish ovaries, as demonstrated by the loss of the piRNA enzymes *zili* and *hen1*, which did not severely abolish early germ cells in gonads, but resulted in later loss of abnormally differentiating oocytes [4,6]. These dynamics of piRNA loss-of-function are consistent with *asz1* phenotypes: ***1)*** loss of *asz1* leads to severe loss of germ cell by at least partly by apoptosis, likely by the up-regulation of transposon expression (Fig. 3N, 4E), similar to the *ziwi* mutant phenotype [4], ***2)*** the rare cases of rescue of germ cells from apoptosis in *asz1;tp53* double mutants allows oocyte survival, but reveals defects in oogenesis similar to the milder *zili* and *hen1* phenotypes [4,6]. Despite many efforts, we could only recover partial rescue with low number of *asz1^-/-^;tp53^-/-^* ovaries, which preclude further analyses. Nevertheless, the above phenotypes conclude that Asz1 is essential for oogenesis.

In addition to piRNA loss-of-function as measured by increased transposon expression, we detected obscure morphology of piRNA granule organization (Fig. 7B). In zebrafish, the mAb414 antibody was shown to detect an unknown FG-repeat protein that co-localizes with Vasa to piRNA granules in ovaries and to germ granules in PGCs [13,38]. Using mAb414 labeling in *asz1^-/-^* developing gonads, we found that the mAb414 signal exhibited large coalesced aggregates in the cytoplasm of early pre-meiotic germ cells, instead of its normal perinuclear granule organization in the wt (Fig. 2B, 3I). A similar coalescence of mAb414-positive granules into large aggregates, and/or their ectopic distribution in the cytoplasm was detected in *asz1^-/-^;tp53^-/-^* oocytes (Fig. 5M-N). These observations suggest structural defects of piRNA granules in the absence of Asz1 (Fig. 7B). Interestingly, co-labeling of Vasa with mAb414 exhibited normal Vasa granule localization in the same cells that showed the mAb414 phenotypes above. The nature of RNP complex composition and hierarchy in zebrafish germline granules is still unclear, and it is possible that Asz1 is required for the integrity and/or regulation of some complexes in piRNA granules, but not for others. The identification of the yet unknown FG-repeat protein that is detected by mAb414 in zebrafish germline granules is required for further investigation.

Considering the roles of Asz1 in piRNA granules, as well as the importance of different types RNP granules for germline development, we tested the potential role of Asz1 in Bb granules. Asz1 localizes to the mature Bb in zebrafish and Xenopus [13,20,34], and undergoes concomitant polarization dynamics with Bb formation in zebrafish [13]. However, examining Bb granule components – the Buc protein and the *dazl* mRNA, in *asz1^-/-^;tp53^-/-^* oocytes, showed their normal localization to the forming Bb, as well as to the mature Bb (Fig. 6A-D, S4-5). These results demonstrated that the Asz1 protein is not essential for the localization of major Bb granule components to the Bb, or for major steps in Bb formation (Fig. 7C). Since only a few Bb proteins have been identified so far, including Buc [20], Macf1 [50], Tdrd6a [51], and Rbpms [22], it is possible that ***1)*** Asz1 is required for Bb structure/function downstream of Buc and *dazl*, and/or ***2)*** Asz1 affects other components in ways that are not detected by Buc and *dazl* labeling, and/or ***3)*** Asz1 functions redundantly in the Bb. It is also possible that the Asz1 protein does not function in the Bb, but uses the Bb as means for localization in the oocyte [19], towards potential maternal functions in the future embryo (see below). The exclusive development of *asz1* mutants as sterile males, precludes further analyses of potential functions of the Bb and/or its components later in oogenesis or embryogenesis.

The differential necessity for Asz1 in piRNA granules versus Bb granules sheds interesting light on RNP complexes. The full repertoire of proteins and transcripts in both piRNA and Bb granules is far from being identified. Moreover, both homotypic and heterotypic complexes have been demonstrated to form in hierarchy in polar granules in the Drosophila germline [52–56]. However, whether RNP interactions form heterotypic or homotypic complexes or both, and their hierarchy of formation is not understood in zebrafish germline granules. A few proteins show overlapping and distinct localization and function between granule types. For example, Buc localizes to both germ granules in PGCs [3,57,58] and Bb granules [13,57] and is essential for both [20,57,59]. Tdrd6a localizes to all three granule types [51], is essential for PGCs, dispensable in the piRNA pathway, and modifies Buc aggregation and Bb morphology [51]. Here, we show that Asz1 localizes to both piRNA and Bb granules, but is only essential for piRNA function, adding an important piece to the puzzle of zebrafish germline granule organization.

Finally, we show that zygotic Asz1 is dispensable for embryonic PGC specification and migration to the gonad, by monitoring Vasa-positive gem cells in embryos and larvae (Fig. 4). It is possible that some defects in *asz1* mutant PGCs are not detected in this assay. However, the number of PGCs we detected at 24 hpf is consistent with the literature [48], ruling out specification defects. Further, migration at both 24 hpf and 7 dpf was complete, without lagging or mis-localizing PGCs. It is thus likely that zygotic Asz1 is not a major functional regulator of PGC specification and migration in zebrafish. Nevertheless, in addition to the zygotic expression of *asz1* as detected in embryos at 8 and 24 hpf, our expression studies showed that *asz1* transcripts are detected in embryos at 1.5 and 5hpf (Fig. S1A), which is likely due to maternal deposition. It is possible that maternal Asz1 is required for germ plasm regulation and early PGC specification and/or migration. However, the lack of sexually mature *asz1^-/-^* females precludes further analyses of potential maternal Asz1 functions in PGCs.

Asz1 functions are conserved in mammals. Asz1 *gain-of-function* was shown to promote PGC differentiation form mouse and human embryonic stem cells, and to enhance the expression of PGC markers [66], while its *loss-of-function* in mice embryos down-regulated the expression of those markers [66]. In mice, Asz1 is essential in piRNA regulation in the male [29–32], but despite expression in oocytes [28], is dispensable for female fertility [29]. In humans, asz1 is specifically expressed in adult testes according to the human genome atlas, but its functions, in particular potential functions in developing ovaries, have not been determined. Our studies in zebrafish provide new insight into Asz1 functions that can be directly relevant for human reproduction.

In summary, our work demonstrates the function of the Asz1 protein in the zebrafish germline, and sheds new light on the piRNA pathway, the Bb, and germline RNP granules. By identifying a new regulator of germ cell and gonad development in zebrafish and deciphering its mechanistic requirements, our study contributes to advancing our understanding of fertility and reproduction, as well as generally to the field of RNP granules.

## Material and Methods

### Ethics statement

All animal experiments were supervised by the Hebrew University Authority for Biological Models, according to IACUC and accredited by AAALAC. All experiments were appropriately approved under ethics requests MD-18-15600-2.

### Fish lines and gonad collections

Gonads were collected from juvenile and adult fish at indicated developmental stages. Gonad collection was done as in [13,60]. Briefly, fish were cut along the ventral midline and the lateral body wall was removed. For juvenile fish, the head and tail were removed and, the trunk pieces, with the exposed abdomen containing the ovaries were fixed in 4% PFA at 4°C overnight with nutation. Trunks were then washed in PBS and ovaries were finely dissected in cold PBS. For adult fish, gonads were directly removed and fixed. Gonads were washed in PBS and then either stored in PBS at 4°C in the dark or dehydrated and stored in 100% MeOH at -20°C in the dark. Gonads were imaged using a Leica S9i stereomicroscope and camera.

Fish lines used in this research are: TU wild type, *asz1^huj102^* (this work, see below), and *tp53^M214K^* [61].

### Generation of *asz1* mutant fish by Crispr/Cas9

1-cell stage embryos were injected with 1nl of gRNA duplex (crRNA:tracrRNA, 250 ng/µL) and Cas9 protein (PNA-Bio CP01-50; 500ng/µL), as described in [62]. The crRNA and tracrRNA were manufactured by IDT. The *asz1* locus was targeted by two gRNA sequences, designed by the CRISPRscan software [63]. gRNA1 was GTGCGGGAGTTGTTGGATGGTGG (including the PAM sequence) and targeted the middle of Ankyrin repeats domain in exon 2. gRNA2 was AGGGAAGATGGCAGCCGACATGG (including the PAM sequence) and targeted the end of the Ankyrin repeats domain in exon 6.

F0 embryos were raised to adulthood and crossed with wt fish to screen for germline transmission. To identify potential loss-of-function mutations DNA was extracted from F1 fish using the *RAPD* method (see below), followed by genomic PCR amplification using Phusion DNA polymerase, targeting the *asz1* target sequence with primers - Forward: AATCGCAGTGCAGTATGGACA, Reverse: TCCTACTGTGATCTCTTACCGTCA, yielding a 170bp amplicon in the wt. PCR products were analyzed by both Metaphore gel electrophoresis (2% resolution), and by Sanger sequencing (Fig. S1). Positive F1 fish were outcrossed to raise individual heterozygous F2 families, which were then in-crossed to generate F3 fish with Mendelian segregation of *asz1* alleles. From gRNA1, we identified a family with a mutation including a four base pair deletion, generating a premature STOP codon at amino acid 243 (Fig. S1D). We named this allele *asz1^huj102^*. Despite numerous screening efforts in F0 through F3 fish, gRNA2 only produced various mismatch mutations that were not predicted to alter the Asz1 protein sequence and function.

### Genotyping

Fish were anaesthetized in 0.02% of Tricaine (Sigma Aldrich, #A5040) in system water, and fin-clipped, followed by *RAPD* DNA extraction protocol [64]. Briefly, tails are dehydrated in MeOH and lysed and DNA is extracted in in RAPD buffer [64]. For *asz1^huj102^*, genotyping was performed by PCR amplification followed by Metapore gel electrophoresis using the primers - Forward: AATCGCAGTGCAGTATGGACA, Reverse: TCCTACTGTGATCTCTTACCGTCA, yielding a 170bp amplicon in the wt and 166bp in the mutant (Fig. S1). For *tp53*, we performed KASP genotyping (LGC, Teddington, UK), using the following SNP sequence: GATTTACAAACTCTTTTTTTATTTTTATTTTATTATTATTATTATTATTATTATTTACATGAAATTGCCAGAGTATGTGTCTGTCCATCTGTTTAACAGTCACATTTTCCTGTTTTTGCAGCTTGGTGCTGAATGGACAACTGTGCTACTAAACTACATGTGCAATAGCAGCTGCATGGGGGGGA[T/A]GAACCGCAGGCCCATCCTCACAATCATCACTCTGGAGACTCAGGAGTAAGTACTGCATATTTGATTCCTCCTCTTGTGAACTGCTTTTTTAAATTTATTTTTTATTTTTTTGAT.

### RNA extraction and RT-PCT

RNA was extracted from embryos, larvae, juvenile and adult gonads, and adult organs, using TRI-reagent^®^, followed by phenol;chloroform extraction and precipitation, DNAaseI reaction and additional phenol;chloroform extraction and precipitation. cDNA was generated using random hexamer primers and the Superscript IV Reverse Transcriptase kit^®^ as per the manufacturer instructions. cDNA was then used in PCR amplification followed by gel electrophoresis, or by qPCR, using a StepOnePlus Real-Time PCR machine (Applied Biosystems) and SYBRgreen^®^. Primers were:

*Vasa* For 5′-GGTCGTGGAAAGATTGGCCTG-3′,
*Vasa* Rev 5′-CAGCAGCCATTCTTTGAATATCTTC-3′,
*GypsyDR2* For 5′-GAAATCACCTGTGCATTTAC-3′,
*GypsyDR2* Rev 5′-ATGCAGACATTGGGTAAAGC-3′,
*EnSpmN1* For 5′-GATTGGCCATTGTGTTCACATGC,
*EnSpmN1* Rev 5′-GCTGTGACTGTCATAGGTTTACC-3′,
*Ngaro* For 5′-GGGAGCGATCGAGACCTACC,
*Ngaro* Rev 5′-CAATCATATCACGTGCTCCTCTCG-3′,
*Polinton* For 5′-CCTGACAATGTTGTCAGCCTG-3′,
*Polinton* Rev 5′-CATGAAAGCTAAGGGTATAACTCTG-3′,
*DIRS1a* For 5′-GGGTGCGTCACGCTTGC-3′,
*DIRS1a* Rev 5′-GTAACCTCGAACGTTCCCC-3′,
*L1-5* For 5′-GCACAAAGGACAAATTCACTGGAC-3′,
*L1-5* Rev 5′-GTCCACGTTTAGTATTACAGTTGC-3′,
*LTR2* For 5′-GGTGTCGTTAGAATGCCCTTGAC-3′,
*LTR2* Rev 5′-GGTTATACCTGTGGGTCACGTG-3′.

### Fluorescence immunohistochemistry (IHC) and RNA-FISH by HCR

For gonads, IHC was performed as in [13,60]. Briefly, gonads were washed 2 times for 5 minutes (2×5min) in PBT (0.3% Triton X-100 in 1xPBS; if stored in MeOH, gonads were gradually rehydrated first), then washed 4×20min in PBT. Gonads were blocked for 1.5-2 hours (hr) in blocking solution (10% FBS in PBT) at room temperature, and then incubated with primary antibodies in blocking solution at 4°C overnight. Gonads were washed 4×20min in PBT and incubated with secondary antibodies in fresh blocking solution for 2 hr, and were light protected from this step onward. Gonads were washed 4×20min in PBT and then incubated in PBT containing DAPI (1:1000, Molecular Probes), with or without DiOC6 (1:5000, Molecular Probes) for 50 min and washed 2×5min in PBT and 2×5min in PBS. All steps were carried out with nutation. Gonads were transferred into Vectashield (with DAPI, Vector labs). Gonads were finally mounted between two #1.5 coverslips using a 120 μm spacer (Molecular Probes).

For embryos, IHC was performed similarly, except that embryos were always stored in MeOH at -20C, and rehydrated before labeling, followed by five 5 minutes washes in PBT. Embryos were then treated with 5µg Proteinase K for 1 min, re-fixed in 4% PFA for 20min, and washed three times in PBT for 5 minutes, before blocking.

Primary antibodies used were Vasa (1:5000, [39]), mAb414 (1;1000, Abcam), Acetylated tubulin (1:200; Sigma-Aldrich), β-Catenin (1:1000; Sigma-Aldrich), cCaspase3 (1:300, Abcam), Buc (1:400, [19]). Secondary antibodies were used at 1:500 (Alexa-flour, Molecular Probes).

RNA-FISH was performed using the third generation DNA-HCR-FISH technique (Molecular Instruments) [65], as in [13,60] and following the company protocol, except for the hybridization temperature that was optimized for 33°C.

### Confocal Microscopy, Image Acquisition and Processing

Images were acquired on a Zeiss LSM880 confocal microscope using a 40X lens. The acquisition setting was set between samples and experiments to: XY resolution=1104×1104 pixels, 12-bit, 2x sampling averaging, pixel dwell time=0.59sec, zoom=0.8X, pinhole adjusted to 1.1μm of Z thickness, increments between images in stacks were 0.53μm, laser power and gain were set in an antibody-dependent manner to 7-11% and 400-650, respectively, and below saturation condition. Unless otherwise noted, shown images are partial Sum Z-projection. Acquired images were not manipulated in a non-linear manner, and only contrast/brightness were adjusted. All figures were made using Adobe Photoshop CC 2014.

### In vitro fertilization (IVF)

IVF was performed as in [39]. Briefly, sperm were collected from anesthetized males into Hank’s solution and stored on ice until eggs were collected. Sperm from individual males of either wt or *asz1^-/-^* strains, was used to fertilize eggs collected from control wt females. Anesthetized wt females were placed in a dish and squeezed for egg collection. 100-150ul sperm solution was added to the collected eggs and incubated for 20 seconds. 1ml of E3 containing 0.5% fructose was added to activate sperm, gently mixed and incubated for 2 minutes. 2 ml of E3 was added followed by 5 minutes incubation. The dish was then flooded with E3 and placed in a 28°C incubator until examination of fertilization rates.

### Statistical analysis

All statistical analysis and data plotting was performed using the GraphPad Prism 7 software. Data sets were tested with two-tailed unpaired *t*-test, unless otherwise indicated. *p*-values were: *<0.05, **<0.01, ***<0.001, ****<0.0001, ns=not significant (>0.05).

## Supporting information

All Supplemental Figures 1-5

## Acknowledgments

We thank Holger Knout for kindly sharing the Vasa antibody, and for Maayan Visuals for the preparation of the schematic image in Fig. 7.

## Author contributions

Conceptualization and Writing: AA, YB, YME

Methodology: AA, GS, YME

Investigation and Visualization: AA, GS, YB, YME

Funding acquisition: YME

Supervision: YME

## Data and materials availability

All data are available in the main text or the supplementary materials.

## Supplementary Materials

### Supplementary Figure legends

**Figure S1.**
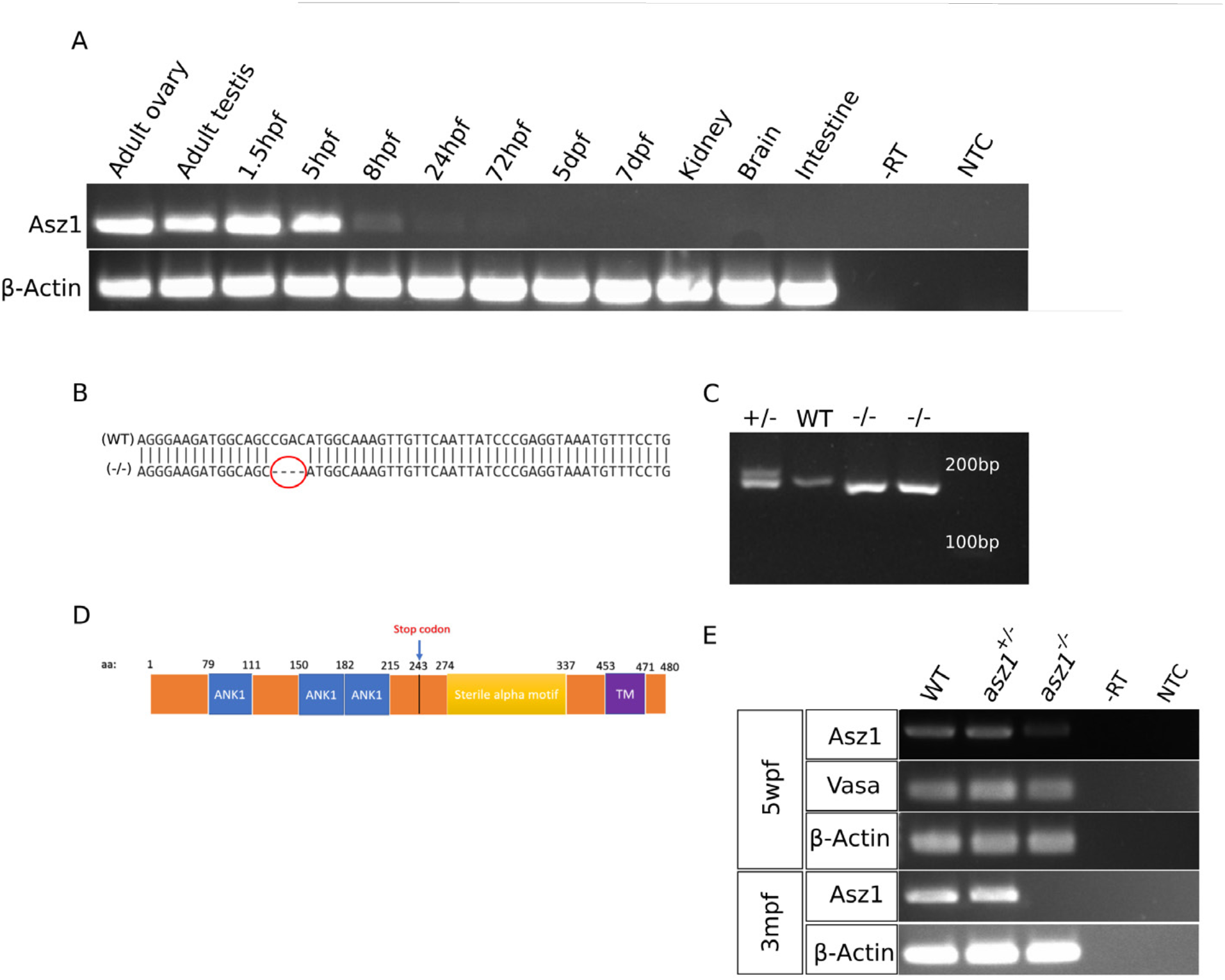
Asz1 expression and the *asz1^huj102^* mutant. **A.** RT-PCR analysis showing specific expression of *asz1* in ovaries and testes, with maternal deposition in early embryos, and detection of likely zygotic transcripts in 8 and 24 hpf embryos. -RT – no reverse transcriptase control; NTC – no DNA template control. **B.** wt and *asz1^huj102^* genomic sequence (from Sanger sequencing) showing a 4 bp deletion in *asz1^huj102^*, leading to premature Stop codon at amino acid position 243 (D). **C.** PCR analysis of genomic DNA from wt, *asz1^+/huj102^*and *asz1^huj102/huji102^*fish, followed by Metapore gel electrophoresis (with 2% electrophoresis resolution), showing distinct bands of the *asz1* locus in the three genotypes (individual fish per genotype; two homozygous fish are shown). Amplicons are 170 and 166 bp in the wt and *asz1^huj102^* alleles, respectively. **D.** A scheme of the zebrafish Asz1 protein, showing the Ankyrin repeats, Sterile alpha motif, and transmembrane domain. The top arrow indicates the position of the premature STOP codon at amino acid position 243 in the *asz1^huj102^*allele. **E.** RT-PCR analysis of asz1 expression in wt, *asz1^+/huj102^* and *asz1^huj102/huji102^* gonads at 5 wpf (n=50 gonads per genotype) and 3 mpf (n=6 gonads per genotype). *β-act* expression serves as loading control. *Vasa* expression marks the presence of germ cells at 5 wpf. *asz1* transcripts are not detected in 3 mpf gonads. At 5 wpf, *asz1* transcripts are only very weakly detected, despite vasa expression, demonstrating the likely decay of the *asz1* mRNA by nonsense mediated decay. -RT – no reverse transcriptase control; NTC – no DNA template control.

**Figure S2.**
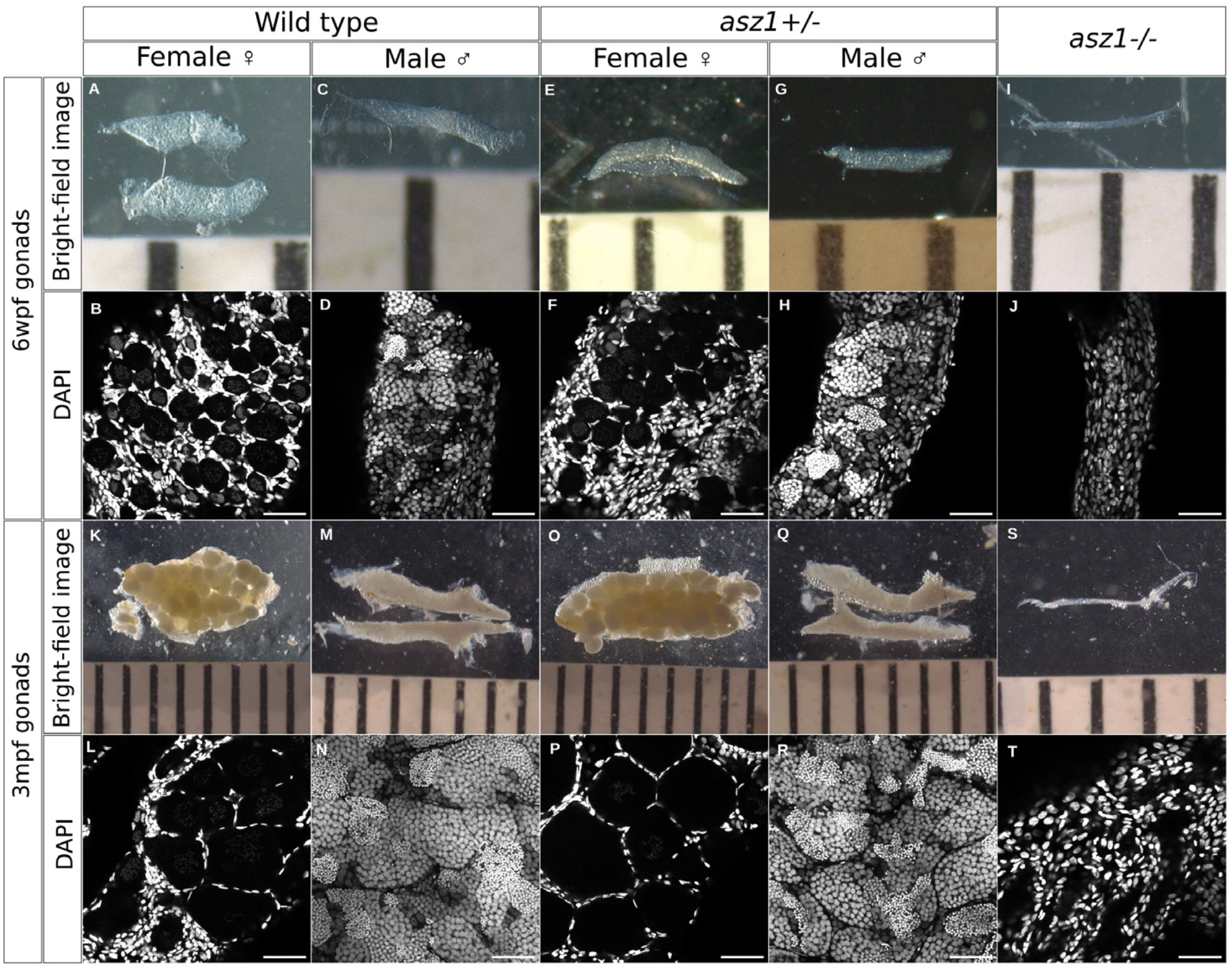
Loss of Asz1 leads to severe gonad developmental defect and germ cell loss. **A, C, E, G, I.** Representative brightfield images of wt, *asz1^+/-^*, or *asz1^-/-^* gonads at 6 wpf. Ruler grades are 1 mm. **B, D, F, H, J.** Confocal images of the same gonads from the brightfield images above, labeled for DAPI (greyscale). Wt and *asz1^+/-^* gonads exhibited normal developing oocytes and early spermatocytes in ovaries and testes as indicated, as well as oogonia and spermatogonia as generally detected by germ and somatic cell nuclear morphology (DAPI). *asz1^-/-^* gonads were much thinner with no clear detection of presumptive germ cells. Scale bars are 50 μm. **K, M, O, Q, S.** Representative brightfield images of adult wt, *asz1^+/-^*, or *asz1^-/-^* gonads at 3 mpf. Ruler grades are 1 mm. **L, N, P, R, T.** Confocal images of the same adult gonads from the brightfield images above, labeled for DAPI (greyscale). Wt and *asz1^+/-^* gonads exhibited normal ovarian and testes morphology as generally detected by DAPI (oocyte show weaker DAPI signal than their surrounding follicle cells), while *asz1^-/-^*gonads were much thinner with no clear detection of presumptive germ cells, and exhibited gaps in the tissue. Scale bars are 50 μm.

**Figure S3.**
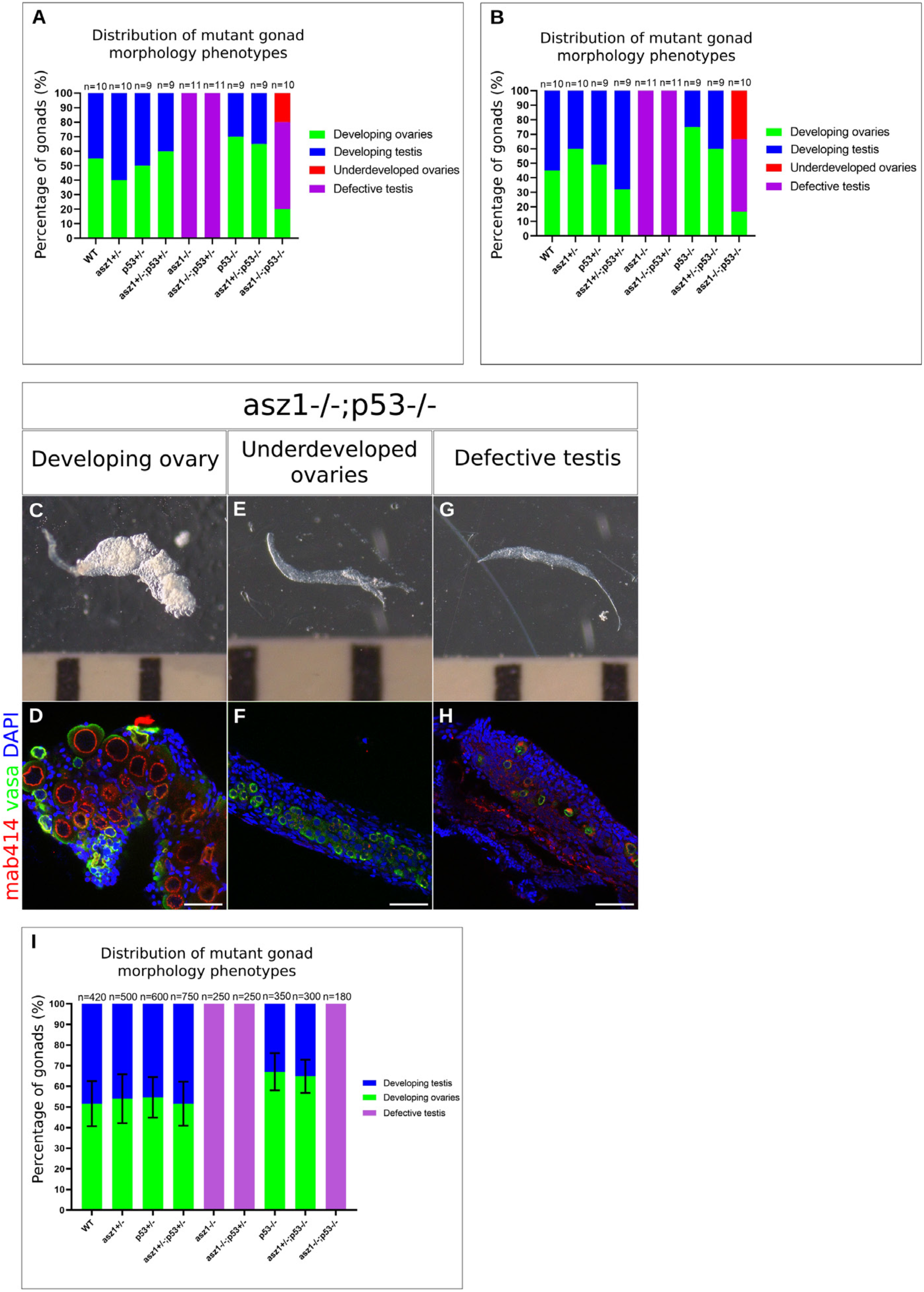
Partial rescue of ovary fate in *asz1;tp53* double mutant fish. **A-B.** The distribution of gonad categories (shown in C-H) per genotype are plotted for the two crosses that yielded ovaries. A and B represent independent crosses. N=number of gonads. **C-H.** Representative images of the gonad categories in *asz1^-/-^;tp53^-/-^* fish. Top panels are brightfield images of the gonads in the bottom panels which are labeled with Vasa (green), mAb414 (red), and DAPI (blue). Scale bars are 50 μm. Developing ovaries are thicker, and contain oogonia and differentiating oocytes. Underdeveloped ovaries are thinner and contain oogonia with no further progressing oocytes. Defective testes are gonads that exhibit the *asz1^-/-^*phenotypes, being very thin and containing very few germ cells with abnormal mAb414 signal. These categories are consistent with our previous description of the morphology of normal and defective juvenile ovaries and testes [39] **I.** The distribution of gonad categories in all crosses that did not yield ovaries in *asz1^-/-^;tp53^-/-^*fish. A total of 17 rounds of in-crosses between 40 *asz1^+/-^;tp53^+/-^*double heterozygous individual fish were performed. In two crosses (panels A-B) a total of 6 *asz1^-/-^;tp53^-/-^* ovaries were obtained. In the remaining 15 crosses (panel I), all 180 *asz1^-/-^;tp53^-/-^*gonads were defective testes. n=number of gonads. Bars are mean ± SD between independent crosses.

**Figure S4.**
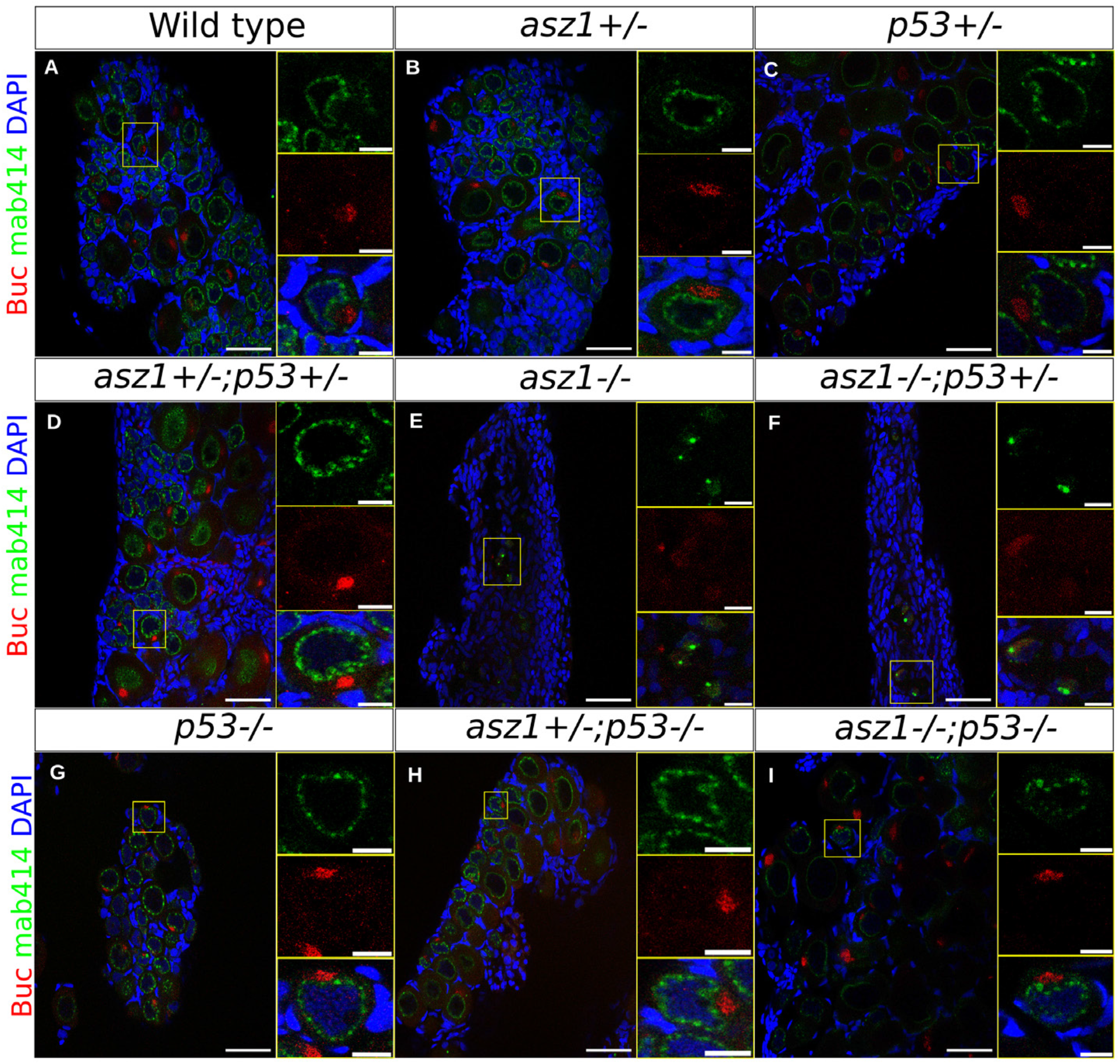
Supporting information for Fig. 6A. Overview of gonads of all *asz1;tp53* genotypes from the experiment in Fig. 6A, labeled for Buc (red), mAb414 (green), and DAPI (blue). Right panels are single and merged channel zoom-in magnifications of the yellow boxes in the overview panels. Scale bars are 50 μm and 10 μm in zoomed out and inset magnification images, respectively. All gonads (ovaries), show normal Buc localization in the forming Bb, except for the testes in *asz1^-/-^*, and *asz1^-/-^;tp53^+/-^*. n=3-6 gonad per genotype, except 2 *asz1^-/-^;tp53^-/-^* ovaries.

**Figure S5.**
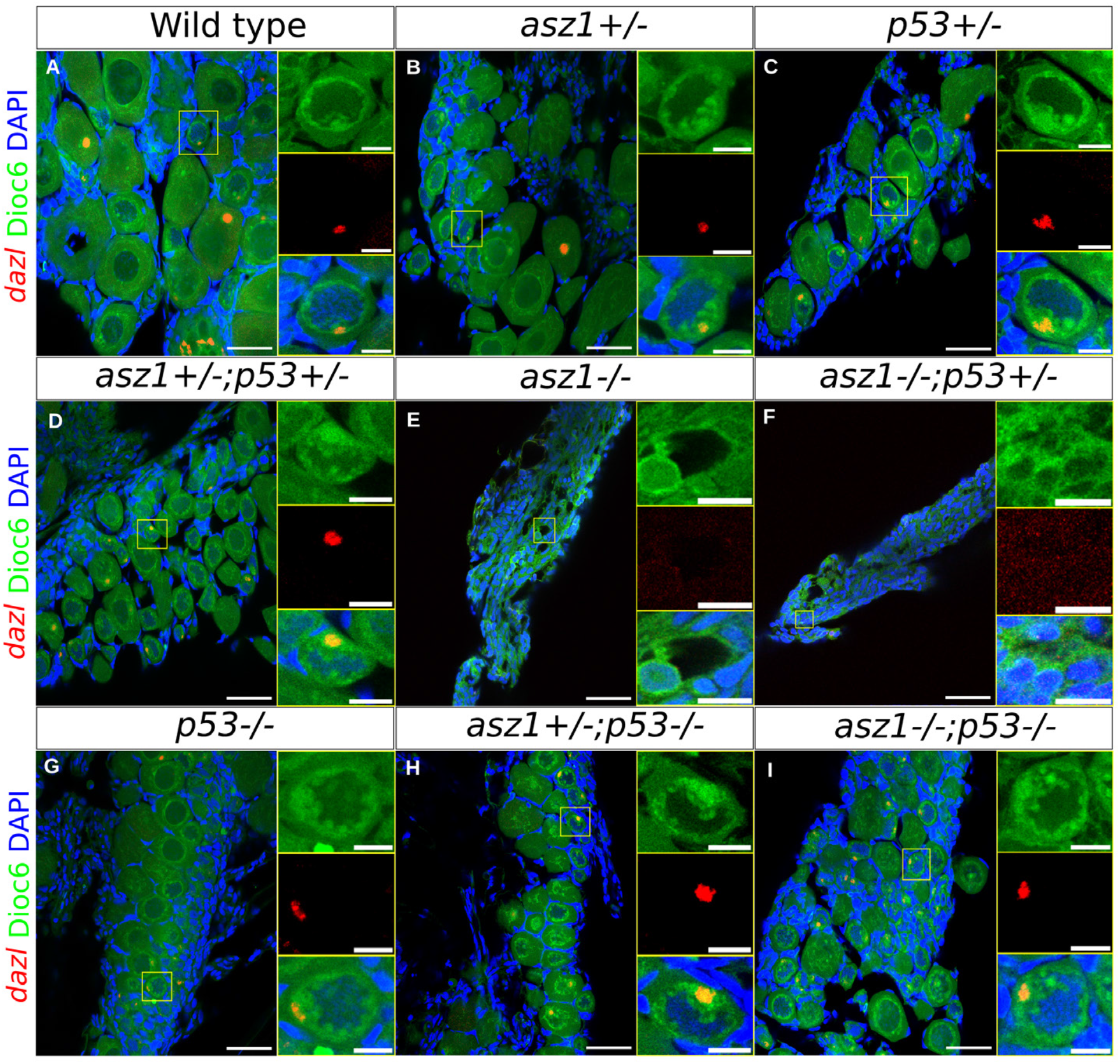
Supporting information for Fig. 6B. Overview of gonads of all *asz1;tp53* genotypes from the experiment in Fig. 6B, labeled for *dazl* (red), DiOC6 (green), and DAPI (blue). Right panels are single and merged channel zoom-in magnifications of the yellow boxes in the overview panels. Scale bars are 50 μm and 10 μm in zoomed out and inset magnification images, respectively. All gonads (ovaries), show normal *dazl* localization in the forming Bb, except for the testes in *asz1^-/-^*, and *asz1^-/-^;tp53^+/-^*. n=3-5 gonad per genotype, except 2 *asz1^-/-^;tp53^-/-^* ovaries.

