## Supplementary figures and images for "The piRNA protein Asz1 is essential for germ cell and gonad development in zebrafish and exhibits differential necessities in distinct types of RNP granules"

### All Supplemental Figures 1-5

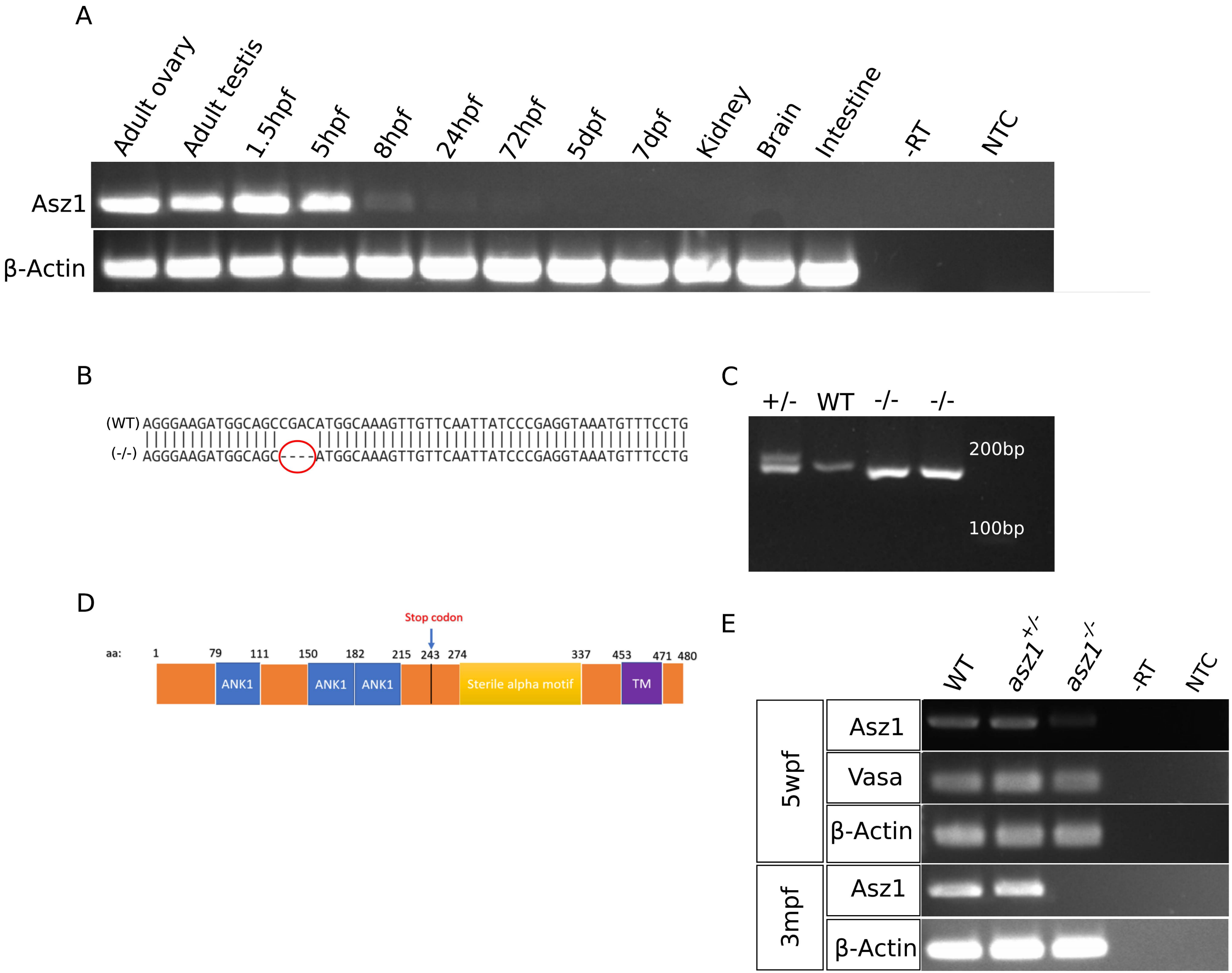

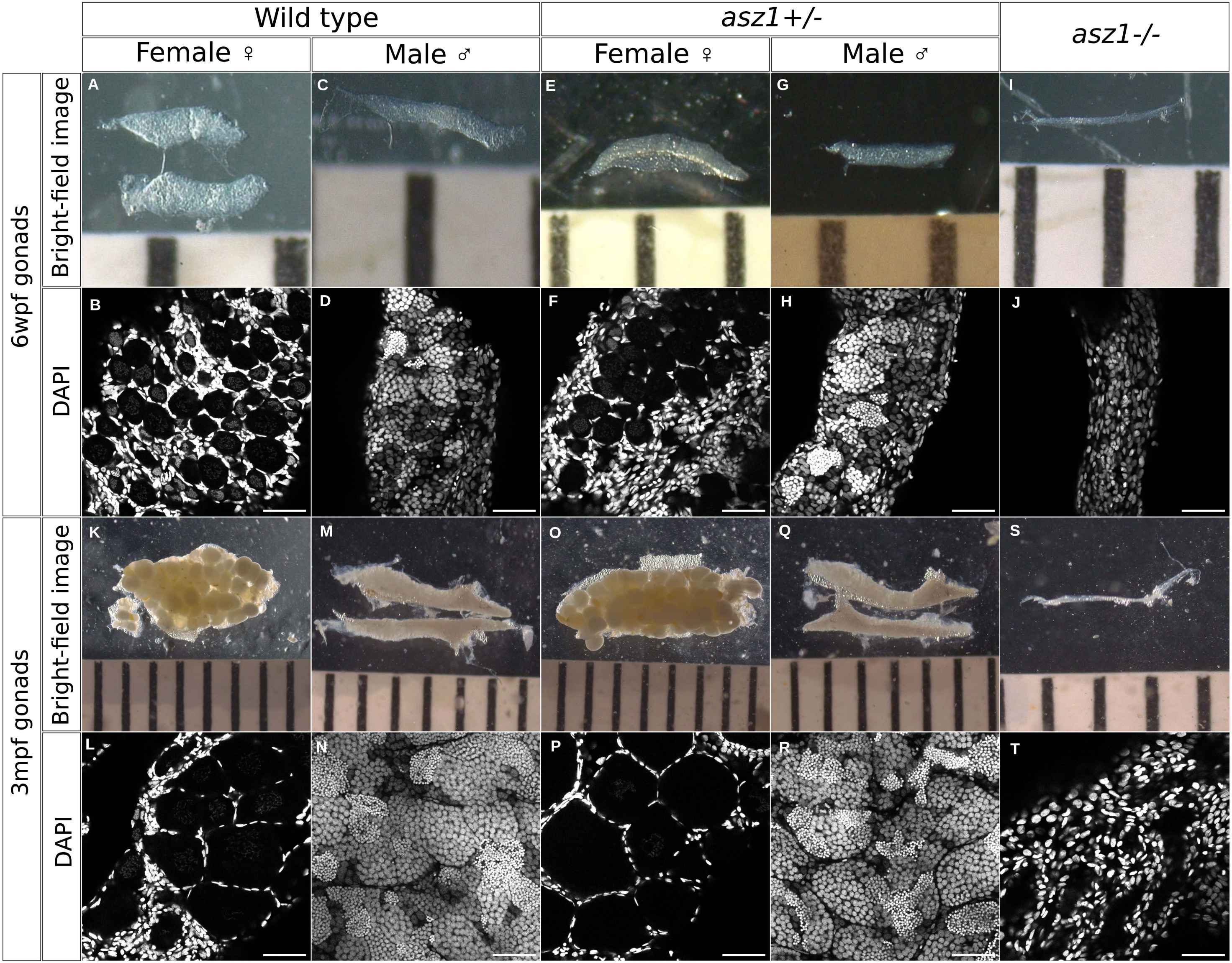

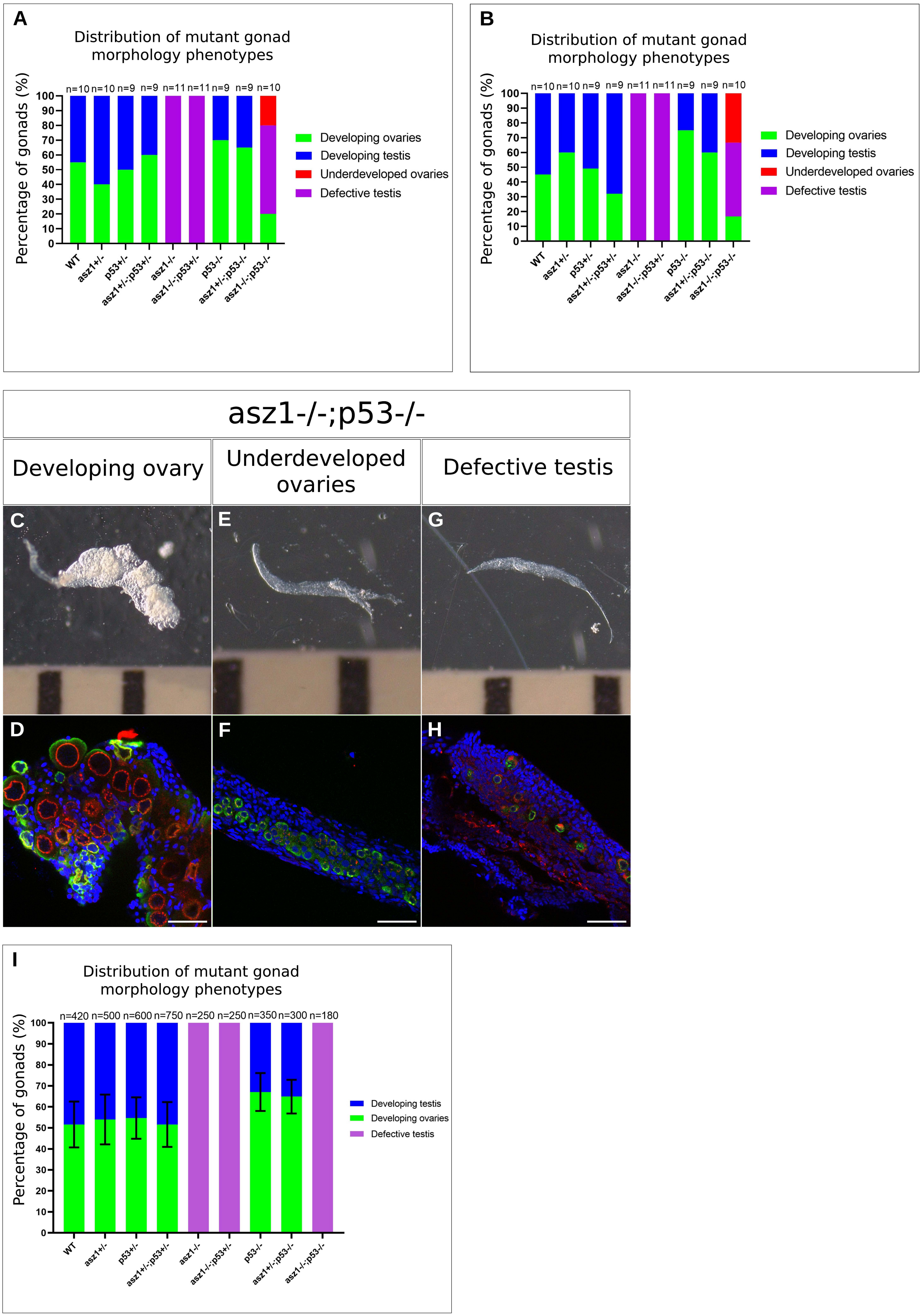

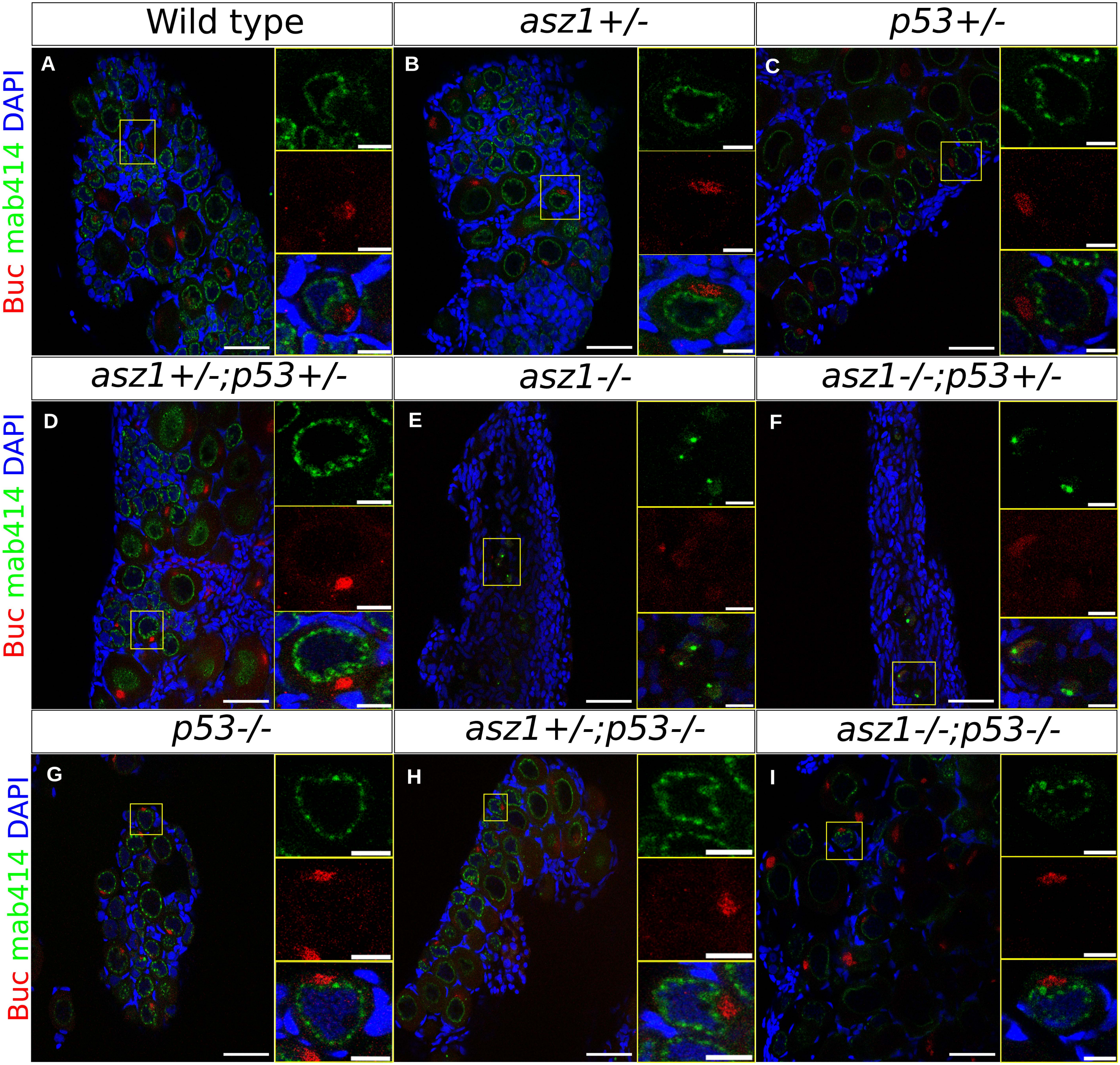

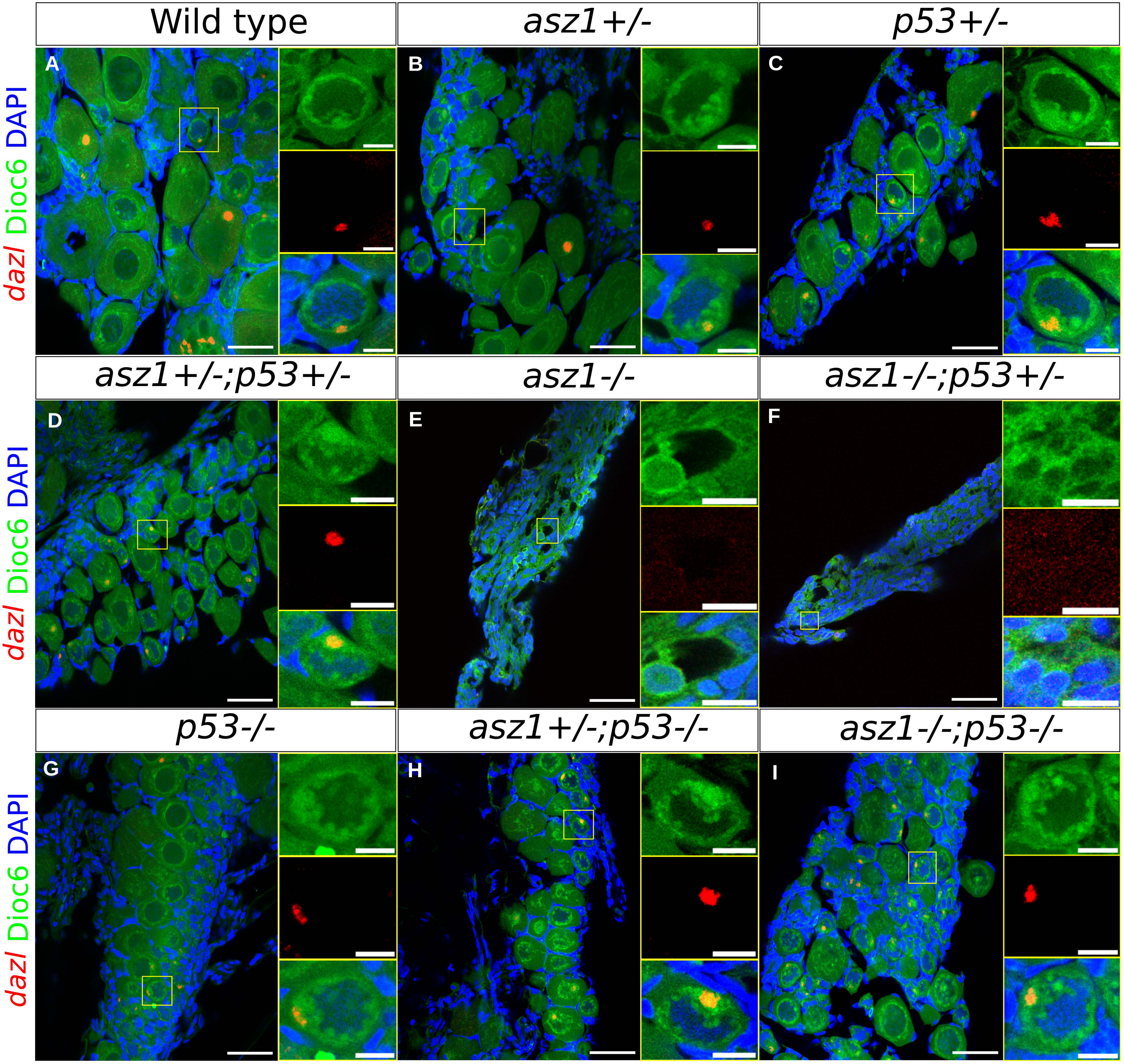
